# GpoA Glutathione Peroxidase Links Oxidative Stress Response, Antibiotic Persistence, and Virulence in *Streptococcus pneumoniae*

**DOI:** 10.64898/2026.01.29.699744

**Authors:** V.E. Zappia, M. Hernandez-Morfa, L. Raya-Plasencia, N.B. Olivero, P.R. Cortes, J. Echenique

**Author notes:** Correspondence: CIBICI (CONICET), Departamento de Bioquímica Clínica, Facultad de Ciencias Quimicas, Universidad Nacional de Córdoba. Medina Allende esq. Haya de la Torre, Ciudad Universitaria. X5000HUA Cordoba, ARGENTINA. (ext. 3143). Department of Host-Microbe Interactions, St. Jude Children’s Research Hospital, Memphis, TN, USA.

## Abstract

Glutathione (GSH) peroxidases are conserved enzymes found in prokaryotic and eukaryotic organisms that reduce H2O_2_ to protect cells from oxidative stress damage. In this study, we identified and characterized the GpoA glutathione peroxidase of *S. pneumoniae*, one of the most important human bacterial pathogens. This bacterium is unable to synthesize endogenous GSH as a cofactor for this enzyme but acquires GSH from host tissues via the GshT transporter. We demonstrated that recombinant GpoA exhibits GSH peroxidase activity and that the Cys^36^ residue is essential for this function. The *gpoA* transcripts, as well as *tpxD* and *ahpD*, which encode the TpxD thiol peroxidase and the AhpD alkylhydroperoxidase, respectively, were upregulated when pneumococci were exposed to H_2_O_2_. The *ΔgpoA, ΔtpxD, ΔahpD*, and *ΔgshT* mutants exhibited increased susceptibility to H_2_O_2_, and also impaired intracellular survival in pneumocytes, macrophages, and neutrophils compared to the wild-type strain. These findings indicate that GpoA, TpxD, and AhpD constitute a robust H_2_O_2_ detoxification system that depends on extracellular GSH uptake. These three peroxidases also contribute to the fluoroquinolone persistence mechanism, which is closely associated with the oxidative stress response in *S. pneumoniae*. Additionally, we investigated the effect of GpoA on virulence in a murine model. The Δ*gpoA* mutant exhibited diminished survival across multiple organs relative to the wild-type strain, suggesting that GpoA contributes to pneumococcal pathogenesis. The molecular, biochemical, and functional analysis of GpoA elucidates an effective bacterial mechanism that incorporates the TpxD and AhpD peroxidases alongside the GshT transporter, promoting extracellular and intracellular survival under oxidative stress conditions.

## 1. Introduction

*Streptococcus pneumoniae* is a Gram-positive bacterium that typically colonizes the human nasopharynx. However, under certain yet poorly understood conditions, it can invade host tissues and transition into a significant pathogen. This microorganism is responsible for a broad spectrum of diseases, ranging from mild infections such as otitis media and sinusitis to severe conditions including meningitis, pneumonia, and bacteremia [1].

In spite of the availability of vaccines and antibiotics, *S. pneumoniae* remains a significant public health concern, with reports indicating it is responsible for over 1.2 million deaths annually, primarily affecting young children and the elderly (https://www.cdc.gov/pneumococcal/global.html). Furthermore, the increasing antibiotic resistance of this pathogen poses substantial challenges to the treatment of invasive pneumococcal diseases. For this reason, the World Health Organization has designated *S. pneumoniae* as a medium-priority pathogen for research and development efforts [2].

During the infectious process, *S. pneumoniae* adapts to varying oxygen levels encountered in different host environments. In the nasopharynx, bacteria are exposed to approximately 20% oxygen, akin to atmospheric levels. Within host cells and the tracheal-respiratory interface, oxygen availability decreases to about 5%, reflecting microaerophilic conditions. In contrast, in severe infections such as bacteremia and meningitis, the pneumococcus must endure completely anaerobic conditions, with 0% oxygen present in the bloodstream and cerebrospinal fluid. Although this pathogen lacks catalase, which is common in prokaryotic and eukaryotic cells to combat oxidative stress, the pneumococcus relies on a unique set of proteins to manage its oxidative stress response (OSR)[3], an specialized mechanism allow it to tolerate oxidative stress and thrive in these diverse oxygen environments. Among the proteins involved in the OSR, the TpxD thiol peroxidase [4, 5] and the SodA superoxide dismutase [5, 6] play key roles; however, many other proteins remain poorly studied or unidentified, leaving significant gaps in our understanding of their specific roles in oxidative stress tolerance and pathogenesis [3].

We have reported that SodA and TpxD are regulated by the StkP-ComE signal transduction pathway and that are essential for intracellular survival of the pneumococcus within pneumocytes [7]. In addition, we have demonstrated that oxidative stress is a major factor influencing antibiotic persistence in *S. pneumoniae*. We revealed that fluoroquinolone persistence is triggered by oxidative stress, allowing a subpopulation of pneumococcal cells to survive lethal antibiotic concentrations without acquiring genetic resistance [8]. Proteins involved in ROS detoxification, such as SodA and TpxD, contribute to this persistence mechanism [8]. This phenomenon is particularly relevant in intracellular environments, where host cells generate ROS as an antimicrobial defense.

On the other hand, glutathione (GSH), a potent non-protein antioxidant, is imported by *S. pneumoniae* via a specific GSH transporter known as GshT. This transporter also plays a critical role in the OSR and the pathogenicity of *S. pneumoniae*, as the bacterium lacks the capability to synthesize GSH endogenously [9]. Glutathione peroxidases (GSH-Pxs) constitute a family of antioxidant enzymes primarily present in eukaryotic organisms, which utilize GSH as a cofactor to catalyze the reduction of H_2_O_2_.These enzymes play a crucial role in antioxidant defense mechanisms and are potentially involved in immune responses against pathogenic infections [10]. For instance, GSH-Px1 in mammals is associated to inflammatory signaling pathways [11, 12]. In contrast, the study of GSH-Pxs in prokaryotes remains underexplored, and they have been reported only in *Streptococcus pyogenes*, where a *gsh-px* mutant was associated to an impaired virulence in mice infection models [13]. However, in *Listeria monocytogenes* the *gpoA* mutant showed improved oxidative stress resistance and pathogenicity using a mice model [14]. Nevertheless, the functional role of GSH-Pxs in bacterial response to oxidative stress and host infection, as well as its biochemical characterization, has been poorly characterized.

In eukaryotes, GSH-Px enzymes catalyze the reduction of organic hydroperoxides and H_2_O_2_ into their corresponding alcohols (or water, in the case of H_2_O_2_) using GSH as an electron donor [15, 16]. The question of whether bacterial GSH-Pxs exhibit similar GSH dependence and how they influence oxidative stress tolerance and pathogenicity in bacteria remains largely unanswered. As mentioned above, studies on *L. monocytogenes* provide a foundation for addressing these gaps, highlighting the need to investigate the biological functions of bacterial GSH-Pxs, particularly in oxidative resistance and host-pathogen interactions [14].

Firstly, we achieved bioinformatics analyses of the pneumococcal R6 genome that revealed the presence of a GSH-Px homolog, *gpoA* (encoded by *spr0285*), which shares homology with GpoA from *Streptococcus pyogenes* [13]. Given the high degree of conservation and the established importance of GSH-Px in oxidative stress resistance in both prokaryotic and eukaryotic cells, we hypothesized that GpoA may play a key role in pneumococcal biology.

In the present study, we identified and characterized the GpoA glutathione peroxidase of *S. pneumoniae*. The enzymatic activity of recombinant GpoA was examined in vitro to elucidate its biochemical properties. Furthermore, we assessed the role of GpoA in conferring resistance to oxidative stress, its involvement in ROS detoxification, its influence on bacterial survival within host cells, and its contribution to persistence during fluoroquinolone treatment. Lastly, the significance of GpoA in virulence was demonstrated using a murine infection model.

## 2. Materials and methods

### 2.1 Bacterial strains and growth conditions

The mutagenesis procedure [17], DNA transformation procedures [18], growth conditions [19], and stock preparation of the pneumococcal strains [20] have been previously reported.

### 2.2 Susceptibility to oxidative stress

Bacterial cultures were grown in Brain Heart Infusion (BHI; pH 7.2) at 37°C until they reached an OD_600nm_ of 0.3. Hydrogen peroxide (H_2_O_2_) was then added to a final concentration of 20 mM, and the cultures were incubated at 37°C for 30 minutes. To assess survival rates, serial dilutions were prepared in Todd Hewitt-Yeast Extract (THYE; pH 7.8) and plated onto Brain Hearth Infusion (BHI) agar plates supplemented with 5% sheep blood. After 24 hours of incubation at 37°C, colony counts were performed to quantify surviving bacteria. Survival percentages were calculated by dividing the number of surviving cells after exposure to 20 mM H_2_O_2_ by the initial cell count prior to the stress treatment. Results are presented as mean percentages ± standard deviation (SD) from three independent experiments performed in triplicate.

### 2.3 Antibiotic persistence assay

Pneumococcal strains were cultured in BHI until mid-log phase (OD_600nm_), exposed to 20 mM H_2_O_2_ for 30 min, followed by treatment with 6 mg/mL levofloxacin for 5 hours at 37°C. After exposure to H_2_O_2_, bacterial cells were centrifuged, washed with PBS, and resuspended in fresh culture medium. At different time points, aliquots were collected, serially diluted, and plated on blood agar plates to determine bacterial survival, as described [8]. Levofloxacin persisters (levopersisters) were quantified as CFU/mL, and minimum inhibitory concentrations (MICs) were assessed using the broth microdilution method, following the Clinical and Laboratory Standards Institute (CLSI) guidelines [21].

### 2.4 Cell lines and culture conditions

The RAW 264.7 (ATCC TIB-61™) and A549 (human lung epithelial carcinoma, pneumocytes type II; ATCC CCL-185™) cell lines were cultured at 37^°^C, 5% CO_2_ in DMEM with 4.5 g/l of glucose, 1% penicillin/streptomycin (P/S) and 10% of heat-inactivated fetal bovine serum (FBS) (Gibco BRL, Gaithersburg, Md.). The undifferentiated human monomyelocyte PLB-985 and *gp91*^*phox*^ KO-PLB-985 (PLB-985-KO) cells [22, 23] were cultured in a complete medium RPMI 1640, supplemented with 1% P/S, and 10 % heat-inactivated FBS. All cell lines were regularly tested for *Acholeplasma, Ureaplasma* and *Mycoplasma* contamination as described [24].

### 2.5 Neutrophil differentiation by DMSO treatment

The differentiation of PLB-985 and PLB-985-KO cells into the neutrophil-like phenotype was performed as reported [22, 23]. The differentiation of PLB-985 and PLB-985-KO to neutrophils was confirmed by determination of CD11b, a cell surface marker, using anti-human CD11b monoclonal antibody (ICRF44; eBioscience) and anti-mouse/Alexa Fluor 647 (BioLegend), as reported [25]. The fluorescence was analyzed by flow cytometry (Attune NxT, ThermoFisher Scientific).

### 2.6 Intracellular survival assays

The intracellular survival assay of pneumococci was conducted as described previously [8, 25]. Briefly, approximately 1.5×10^5^ host cells (A549, RAW 264.7, PBL-985, and PBL-985 *nox2*-KO cell lines) were seeded per well in 12-well plates and cultured in DMEM + 5% FBS at 37°C and 5% CO2 for 24 h until reaching ∼95% confluence, equivalent to approximately 3 × 10^5^ cells. Concurrently, pneumococci were grown in BHI to mid-log phase (OD_600nm_ ∼0.3), and then resuspended in DMEM supplemented with 5% FBS.

The infection of cell monolayers was performed using a multiplicity of infection (MOI) of 30:1 (9 × 10^6^ bacteria: 3 × 10^5^ cells). A549 cells were incubated with pneumococci for 3 hours at 37°C and 5% CO2 to allow bacterial internalization, while the other cell lines were incubated for only 1 h with bacteria. Non-internalized bacteria were removed by discarding the supernatant and replacing it with fresh medium containing gentamicin sulfate (200 μg/mL; US Biological G2030) to kill extracellular bacteria. After 30 minutes of antibiotic treatment at 37°C, the cells were washed three times with PBS. The bacterial survival recorded immediately following the antibiotic treatment was defined as time zero (t = 0).

For the time course of bacterial survival, pneumococci-infected A549 cells were maintained in DMEM + 5% FBS and incubated for an additional 2 hours at 37°C and 5% CO2 after the antibiotic treatment. To monitor cell viability, the cells were trypsinized, and apoptosis/necrosis induced by pneumococcal infection was evaluated using flow cytometry with an Annexin V/propidium iodide labeling kit (Invitrogen), showing ∼5% across all time points. To assess intracellular bacterial survival, cells were lysed by centrifugation at 10,000 g for 5 minutes, and the bacterial pellet was resuspended in THYE medium. Serial dilutions of the lysates were plated on BHI agar containing 5% sheep blood and incubated for 16 hours at 37°C to determine the number of internalized bacteria. The bacterial count at t = 0 was defined as 100% survival, and survival percentages at 2 hours were calculated relative to this baseline. The reference data (100% at t = 0) were excluded from the figures.

### 2.7 Determination of levofloxacin persistence of *S. pneumoniae* in host cells

To assess levofloxacin persistence of pneumococci, A549 and RAW 264.7 cells infected with pneumococci (as described above) were maintained in DMEM supplemented with 1% FBS and 6 mg/mL levofloxacin, while PLB-985 and PLB-985 *nox2*-KO infected cells were cultured in RPMI 1640 with 1% FBS, 1.3% DMSO, and 6 mg/mL levofloxacin. Cell cultures were incubated at 37°C with 5% CO2, and samples were collected at different time points. Host cells were lysed by centrifugation at 15,000 *g* for 10 min, and the resulting bacterial pellet was resuspended in BHI medium. Serial dilutions of lysates were plated on blood agar to quantify the number of intracellular bacteria. The bacterial counts obtained at t=0 were set as 100% survival, and the persistence of pneumococci at later time points was expressed as the percentage of surviving levofloxacin-tolerant cells.

### 2.8 qPCR

Bacterial cells were grown in BHI to the midlog phase (OD_600nm_ 0.3) and exposed to 20 mM H_2_O_2_ for 30 min. Then, RNA purification, cDNA synthesis and qPCR were performed as described [7]. Genes were amplified using oligonucleotides (Table S1) and PowerUp SYBR green Master Mix following the manufacturer’s protocol (Applied Biosystem). The *gyrB* gene was used to normalize the expression in *S. pneumoniae* using the Ct method, as described [26].

### 2.9 Protein expression and purification

The *gpoA* gene from the pneumococcal R6 strain was cloned into the pET15b expression vector and transformed into *Escherichia coli* BL21 (DE3) cells (Thermo Fisher Scientist). Protein overexpression was induced by adding 0.5 mM isopropyl β-D-1-thiogalactopyranoside (IPTG) to bacterial cultures grown at 37°C until an OD_600nm_ of 0.6. After induction, these cultures were incubated for 4 h at 25°C to maximize protein solubility. Bacterial cells were harvested by centrifugation at 6,000 *g* for 15 min at 4°C and resuspended in lysis buffer (50 mM Tris-HCl, 300 mM NaCl, pH 8.0). Bacterial lysis was performed by sonication on ice, and the lysates were clarified by centrifugation at 15,000 *g* for 30 min at 4°C. Soluble His-tagged GpoA protein was purified using nickel affinity chromatography following manufacturer’s instructions (Ni-NTA agarose, Qiagen). Proteins were eluted with an imidazole gradient (20–300 mM imidazole in lysis buffer). Fractions containing purified GpoA were analyzed by SDS-PAGE to confirm expression and purity. Protein samples were mixed with Laemmli sample buffer, boiled at 95°C for 5 minutes, and separated on a 12% SDS-polyacrylamide gel. Gels were stained with Coomassie Brilliant Blue R-250.

### 2.10 Determination of the glutathione peroxidase activity

Glutathione peroxidase activity was quantified using the Glutathione Peroxidase Assay Kit (Cat. # MAK437; Sigma-Aldrich, St. Louis, MO, USA). The assay was performed by monitoring the rate of NADPH consumption, measured as the decrease in optical density at 340 nm (OD_340nm_) over a 5-min period using a BioTek microplate reader. Enzyme activity was calculated according to the manufacturer’s definition of one unit and expressed as units per liter (U/L).

### 2.11 Mice infection model

Murine virulence studies were conducted using specific pathogen-free, 10-week-old BALB/c mice (males). Before mutagenesis, the D39 strain had a previous passage in mice, as described [19]. Animals were randomly divided into three experimental groups and inoculated with either the *wt* D39 strain, the *ΔgpoA* mutant, or the isogenic revertant *gpoA*^+^ strain. All infections were performed using a standardized inoculum of 1 × 10^7^ colony-forming units (CFU) suspended in phosphate-buffered saline (PBS) and administered via the intranasal route using 50 µl/nostril. To assess bacterial dissemination and organ colonization, animals were euthanized via cervical dislocation at 24 h post-infection. Target organs (brain, lungs, liver and spleen) were transferred to sterile plastic tubes. Then, tissues were mechanically homogenized by passage through 70 µm cell strainers, and the resulting homogenates were serially diluted in sterile PBS. Aliquots of the serial dilutions were plated onto blood agar plates and incubated overnight at 37°C. Bacterial burden was subsequently quantified by manual colony counting, with results expressed as CFU per organ.

## 3. Results

### 3.1 The GpoA glutathione peroxidase is part of the H_2_O_2_ detoxification system and contributes to the oxidative stress response of *S. pneumoniae*

Considering the lack of catalases in *S. pneumoniae* [3], we decided to study whether *S. pneumoniae* express a H_2_O_2_ detoxification system similar to that reported in human cells, and we searched in the genome of the pneumococcal D39 strain [27] the homologous genes that constitute this mechanism (Fig. 1). Among these genes, we considered *tpxD* (encodes for a thiol peroxidase or thioredoxin) that has been well characterized [4, 5], *ahpD* (encodes for an alkylhydroperoxidase) that has been reported, although is not clear its participation in the OSR of *S. pneumoniae* [28, 29]. Interestingly, we also found *gpoA*, which encodes for a glutathione peroxidase that have not been reported in *S. pneumoniae* so far. The *gpoA* gene presents a 42% identity (58% homology) with the human GPX4 (Fig. 2) [30], indicating that the structure of GpoA is conserved between bacteria and humans. In addition, *gpoA* from *S. pneumoniae* shares 71% identity (82% homology) with *gpoA* from *S. pyogenes* [13]. A phylogenetically tree showed that GpoA present a higher homology with genes found in bacteria that belong to the Phylum *Bacillota* (before known as *Firmicutes*), mainly in the genus *Streptococcus* (Fig S1).

**Fig 1.**
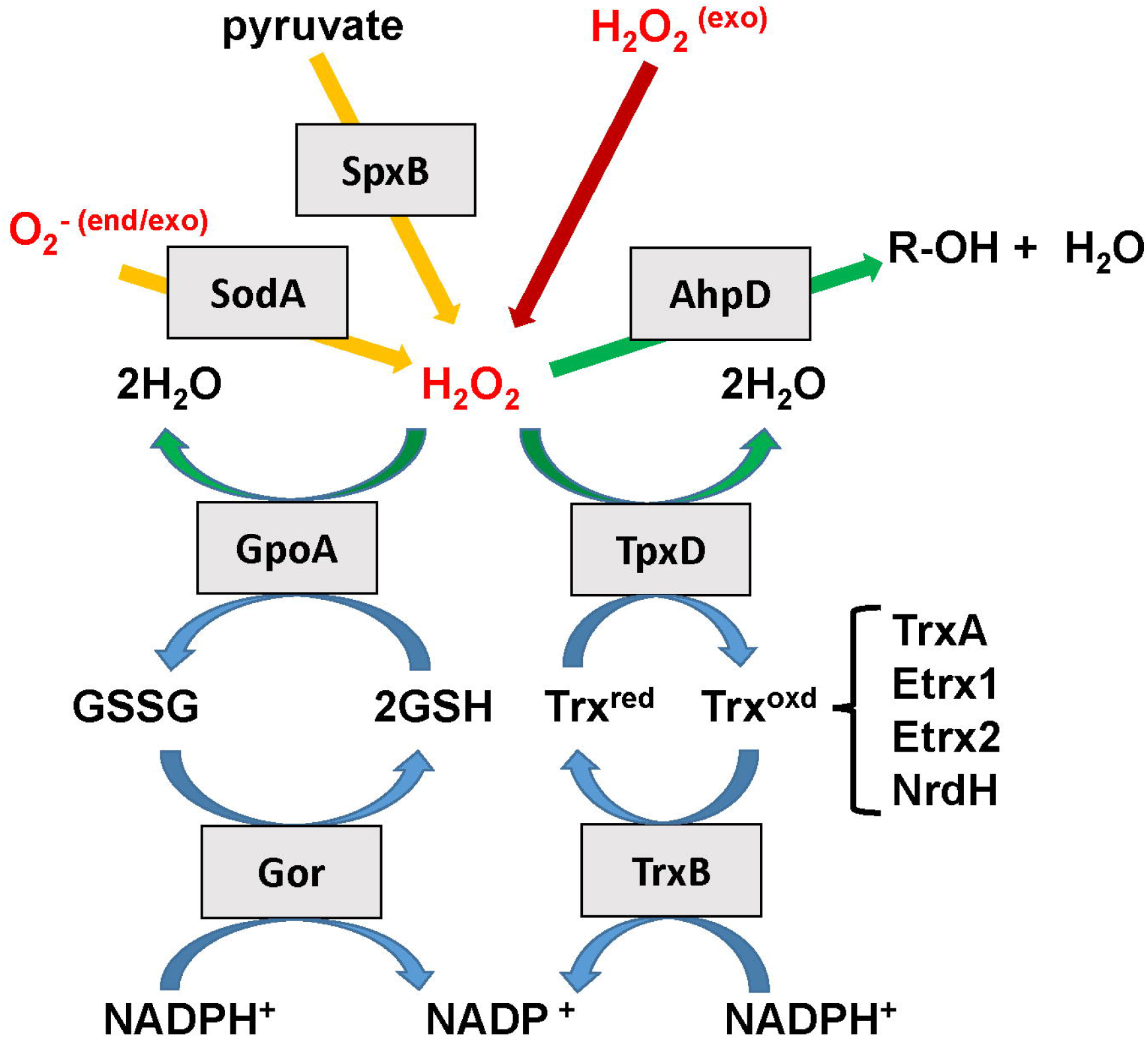
Proposed model illustrating the pathways involved in the production and detoxification of hydrogen peroxide in *S. pneumoniae*. This diagram depicts the oxidative stress response network of *S. pneumoniae*, which must manage both internally produced H_2_O_2_—mainly generated by pyruvate oxidase (SpxB)—and external sources of H_2_O_2_. Superoxide radicals (O2−) are converted into H_2_O_2_ by the superoxide dismutase enzyme SodA. The peroxidases AhpD, GpoA, and TpxD play similar roles in detoxification H_2_O_2_. The detoxification processes dependent on GpoA and TpxD are supported by reductive pathways that recycle cofactors, including glutathione reductase (Gor) for regenerating glutathione (GSH) and the thioredoxin/thioredoxin reductase systems (TrxA/TrxB). Collectively, these pathways maintain redox balance and protect S. *pneumoniae f*rom oxidative stress.

**Fig 2.**
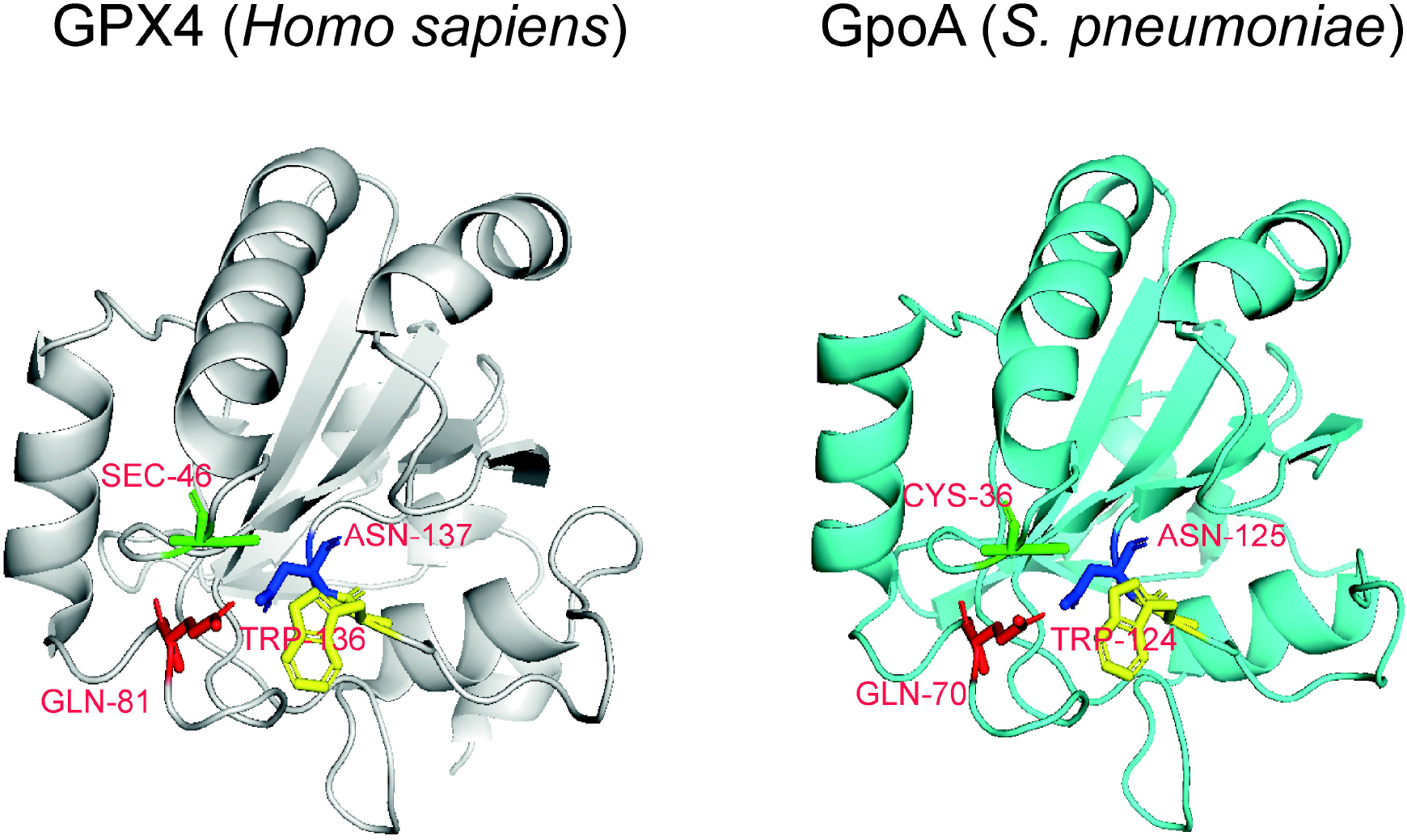
Structural comparison between the human (GPX4) and pneumococcal (GpoA) glutathione peroxidase. Ribbon diagrams of human GPX4 (PDB ID: 2OBI) and pneumococcal GpoA (AlphaFold ID: Q8DR82) were created using PyMOL to emphasize their structural similarities. The amino acids that make up the catalytic tetrad in human GPX4 are highlighted in different colors, along with the corresponding residues predicted to form the catalytic tetrad in GpoA. When the two proteins were superimposed, the root-mean-square deviation (RMSD) was found to be 0.564 Å, demonstrating a strong structural conservation and suggesting that the catalytic framework is preserved between human and bacterial glutathione peroxidases.

To compare the expression of these genes in response to H_2_O_2_, we measured by qPCR the transcriptional level of each one in the pneumococcal R801strain (derivate of the R6 strain) [31] exposed at 20 mM H_2_O_2_ for 30 min, and referred to *gyrB* as internal control [26]. We observed that all the three genes increased significantly their transcript levels after H_2_O_2_ treatment (Fig. 3A), indicating that the increased expression of the *ahpD* and *gpoA* genes are also involved in the OSR of *S. pneumoniae*. This increase was already described for *tpxD* under aerobic cultures compared with bacterial cells cultured under anaerobic conditions [5], but here we report this phenotype under H_2_O_2_ exposition (Fig. 3A). As reference, we also measured the expression of *spxB* (encodes for the pyruvate oxidase) [32, 33] and *sodA* (encodes for a superoxide dismutase) [6] that increased their transcript levels in response to H_2_O_2_, as reported for aerobic conditions [5].

**Fig 3.**
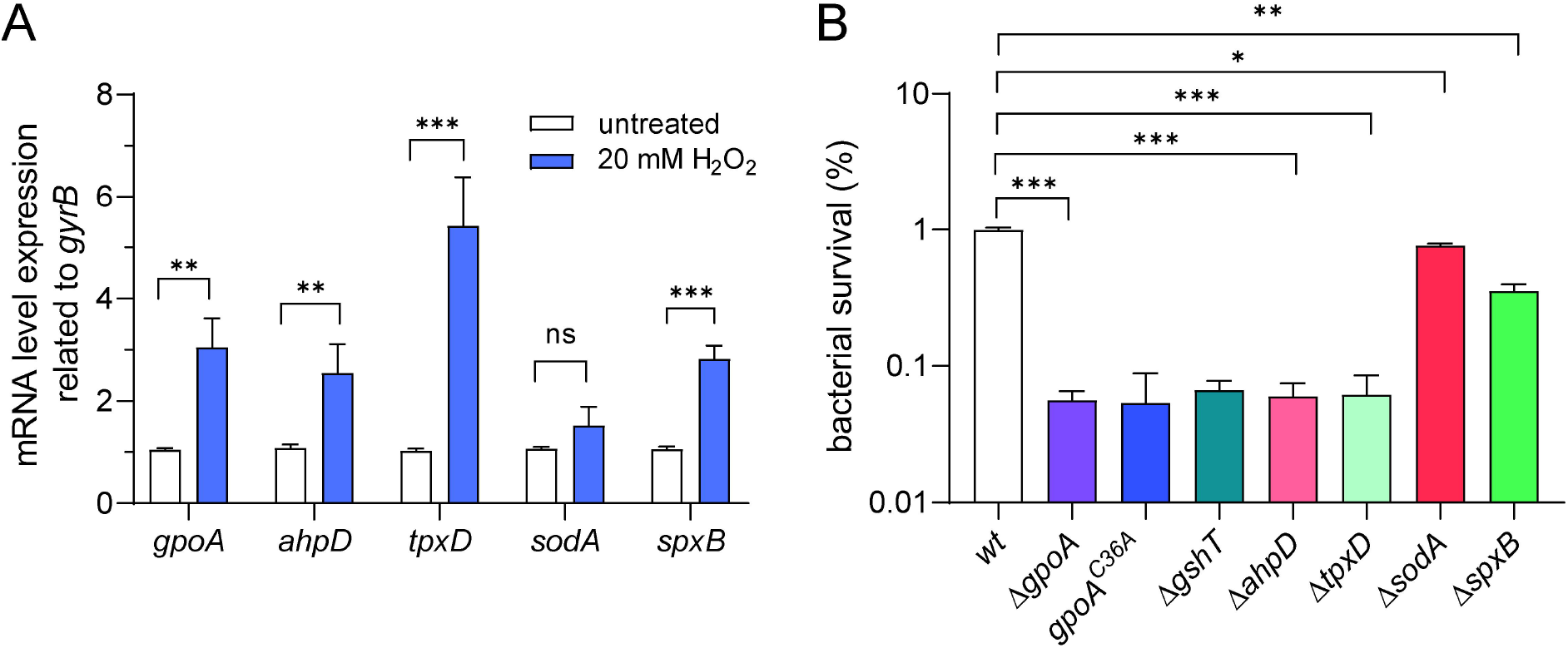
Contribution of pneumococcal peroxidases to the H_2_O_2_ detoxification system. (A) Relative expression of genes involved in the oxidative stress response in the *wt* strain grown under H_2_O_2_-treated and untreated conditions. Gene expression levels were determined by qRT–PCR and normalized to the housekeeping gene *gyrB*, as described in the Materials and Methods section. (B) H_2_O_2_ susceptibility assays for the *wt, ΔgpoA, gpoA*^*C36A*^, *ΔtpxD*, and *ΔahpD* strains. Bacterial cells were exposed to 20 mM H_2_O_2_ for 30 min. Then, pneumococci were washed with PBS and plated on blood agar to estimate CFU/ml. The percentages shown in this panel were calculated from the CFU/ml values presented in Fig. S2. The *ΔsodA* and *ΔspxB* mutants, known to be sensitive to H_2_O_2_, were used as controls. Data in both panels represent the mean ± SEM of at least three replicates. Statistically significant differences were determined using a two-tailed test and are indicated as *P* < 0.05 (*), *P* < 0.01 (**), *P* < 0.001 (***), or *P* < 0.0001 (****). Ns: not significant.

To elucidate the contribution of *gpoA* in the H_2_O_2_ detoxification system of *S. pneumoniae*, and to compare these findings with those reported for *ahpD* [29] and *tpxD* [7], we constructed deletion mutants for each gene in the R801 strain, and we determined H_2_O_2_ susceptibility expressed in percentage of bacterial survival that survived at 20 mM H_2_O_2_ for 30 min [7]. Importantly, the Δ*gpoA*, Δ*ahpD*, and Δ*tpxD* mutants showed a similar H_2_O_2_ susceptibility, representing a 12-fold decrease compared to the wild-type (*wt*) strain (Fig 3B, Fig S2). Previously, it has been reported that the Δ*ahpD* mutant constructed in a R6 genetic background displayed an increased resistance to H_2_O_2_ [29]. This contrasting finding is curious because our R801 strain derivate from the R6 strain. To discard a putative strain dependence of this phenotype, we constructed the *ΔahpD* in a D39 genetic background, the ancestral strain for R6 and R801 strains, and we found the same H_2_O_2_ susceptibility observed in the R801 strain (Fig S3), and this will be discussed later.

Regarding the active site of GSH peroxidases, the selenocysteine localized in position 46 (Sec^46^) showed to be essential for the human GPX4 [30]. In this sense, the orthologous residue found in GpoA is Cys^36^ (Fig. 2), and we proposed that it could be also essential for its peroxidase activity. To test this hypothesis, we constructed the *gpoA*^*Cys36Ala*^ mutant, which demonstrated a higher susceptibility to H_2_O_2_ in reference to the *wt* strain, a similar phenotype displayed by the Δ*gpoA* mutant (Fig 3B, Fig S2). This finding suggested that the Cys^36^ residue is essential for the OSR of *S. pneumoniae*, probably by impairment of the GpoA peroxidase activity. The Δ*gshT* mutant, which lacks the gene that encodes for a GSH transporter [9], confirmed that the intracellular availability of GSH is needed for a typical OSR in *S. pneumoniae* (Fig. 3B). We also tested the Δ*sodA* and Δ*spxB* mutants that have previously shown a lower phenotype than *wt* [6, 32], although these strains displayed a higher H_2_O_2_ resistance than the mentioned mutants, but significantly lower compared with the *wt* strain (Fig 3B, Fig S2). These findings confirmed that GpoA, AhpD and TpxD play a relevant protective role during oxidative stress, potentially enabling the bacterium to persist in hostile environments encountered in different tissues during infection.

### 3.2 GpoA presents GSH-dependent peroxidase activity and requires the Cys^36^ residue

Because GpoA had not been reported so far in *S. pneumoniae*, and has been poorly described in other bacteria [13, 14, 34], we decided to perform a biochemical characterization of this enzyme. To confirm its biological function, firstly we tried to measure the enzymatic activity of GpoA by comparison of total protein extracts obtained from *wt* and Δ*gpoA* strains, but the reactivity of H_2_O_2_ impeded to perform this enzymatic assay. In consequence, we successfully obtained the recombinant GpoA^*wt*^-His_x6_ (wild-type GpoA protein) using the pET15b plasmid expressed in *E. coli* BL21 (DE3), which add the His-tag (His_x6_) at the carboxy-terminal domain of the GpoA protein to facilitate its purification. When total protein extracts were analyzed by SDS-PAGE, a protein overexpression was evident upon IPTG induction, showing a strong band of 18.9 kDa that correspond to the expected molecular weight for GpoA^*wt*^-His_x6_ (Fig. 4A). After the Ni^2^+-NTA affinity purification, the eluted recombinant protein appeared highly enriched, with minimal contamination (Fig. 4A). To assess enzymatic activity of the purified GpoA^*wt*^-His_x6_, we performed a glutathione peroxidase assay, confirming that the purified GpoA retains its expected enzymatic function, that actively catalyzes the GSH oxidation, and that this reaction is dependent on GSH (Fig. 4B).

**Fig 4.**
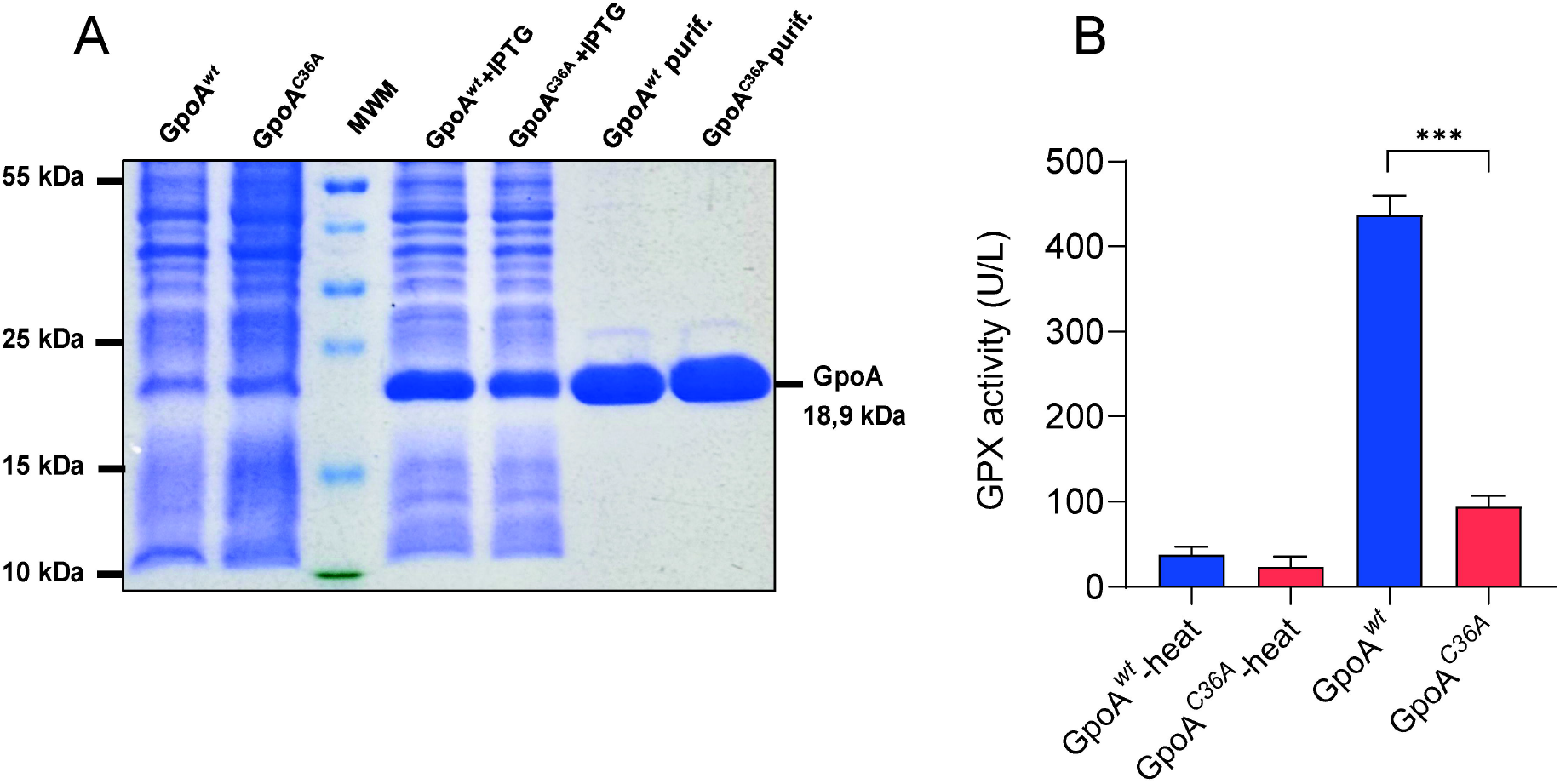
Recombinant GpoA demonstrates glutathione peroxidase activity. (A) SDS– PAGE analysis of total extract proteins of *Escherichia coli* under both non-induced and IPTG-induced conditions, and recombinant proteins purified by columns of Ni-NTA agarose. Both proteins showed bands corresponding to the expected molecular weight of about 18.9 kDa. MWM: molecular weight marker. (B) Glutathione peroxidase activity of purified GpoA and GpoAC36A proteins, measured in units per liter (U/L). The data are presented as the mean ± SEM from at least three independent experiments. Statistically significant differences were assessed using a two-tailed test and are marked with *P <* 0.001 (***).

It has been reported that the catalytic site of the human GPX4 (cytosolic isoform; Uniprot P36969-2) is constituted by a catalytic tetrad that consists of the following residues: selenocysteine (Sec^46^), glutamine (Gln^81^), tryptophan (Trp^136^) and asparagine (Asn^137^) [35]. The comparison with the crystal structure of GPX4 (PDB 6HN3) revealed that a modelated GpoA display orthologous residues that could conform the catalytic site of tis enzyme, including Gln^70^, Trp^124^ and Asn^125^, but Cys^36^ in the place of Sec^46^ (Fig. 2). The selenocysteine present a similar structure than cysteine, but it contains selenium while cysteine contains a sulfur atom. Selenocysteine is an amino acid with a unique genetic encoding system involving a UGA stop codon. To direct the translation of the UGA codon into selenocysteine instead of a stop signal, it is needed a specialized tRNAs for its incorporation into proteins known as Selenocysteine Insertion Sequence (SECIS) [36]. In bacterial proteins, the SECIS (selenocysteine insertion sequence) element is located in the coding region immediately downstream of the UGA codon that encoded for selenocysteine. The Cys^36^ residue in GpoA is encoded by a UGU codon, however, GpoA present a SECIS element identified in silico immediately after this codon that suggest an evolution from selenocysteine to cysteine in *S. pneumoniae* (Fig. S4).

As mentioned, the Sec^46^ residue in GPX4 showed to be essential for the enzymatic activity [35]. In consequence, we hypothesized that Cys^36^ could play the same role in glutathione peroxidase activity of GpoA. To examine the role of this residue from a biochemical perspective, we constructed the recombinant GpoA^Cys36Ala^-His^x6^ mutant using a directed mutagenesis protocol [26]. When we compared the enzymatic activity of both purified recombinant proteins, we observed that GpoA^Cys36Ala^-His_x6_ lost the glutathione peroxidase capacity compared with GpoA^wt^-His_x6_ (Fig 4B). These findings indicated that the Cys^36^ residue is relevant for the glutathione peroxidase activity of *S. pneumoniae*.

### 3.3 GpoA is needed for the intracellular survival of *S. pneumoniae* in host cells

We have demonstrated that the pneumococcus can survive inside pneumocytes, macrophages and neutrophils for several hours [8]. On the other hand, GSH-Pxs are known to be vital components of antioxidant systems in eukaryotes, but their role in bacterial oxidative stress response remains poorly characterized [3]. To deepen assess the impact of *gpoA* deletion on bacterial pathogenesis, we examined the intracellular survival of the *ΔgpoA* mutant in three different host cell types: human pneumocytes (A549; ATCC CCL-185), mouse macrophages (RAW 264.7; ATCC TIB-71), and human neutrophils (PLB-985) [22]. In all three cell models, the *ΔgpoA* mutant exhibited a significantly lower intracellular survival rate compared to the *wt* strain (Fig. 5A), suggesting that GpoA is required for bacterial survival within host cells. This reduced intracellular survival may be attributed to the mutant’s heightened susceptibility to oxidative stress showed before (Fig. 3B), as immune cells generate ROS as a primary defense mechanism against invading pathogens.

**Fig 5.**
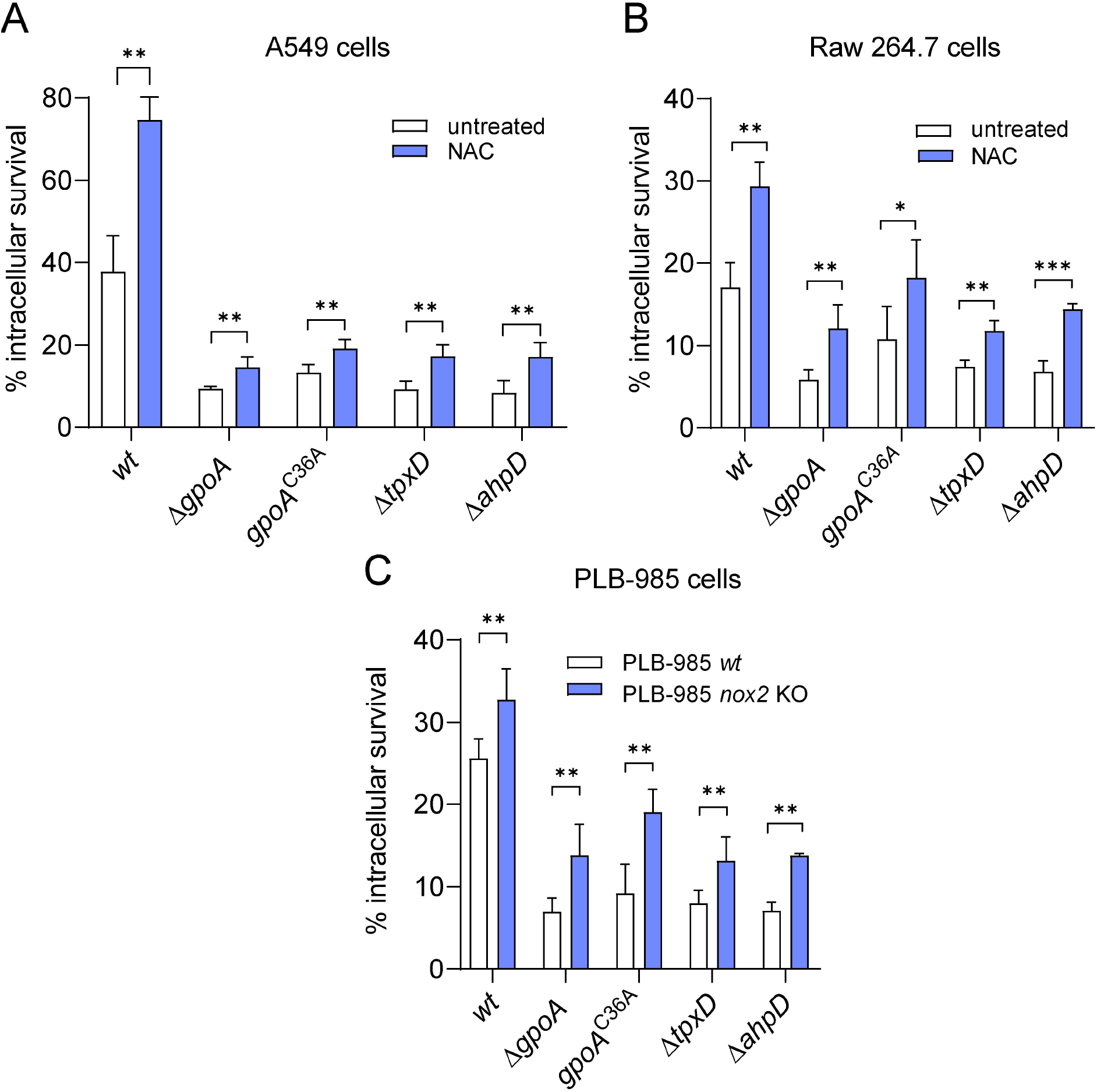
GpoA plays a role in the intracellular survival mechanism of *S. pneumoniae* within host cells. The three panels display the intracellular survival rates of *the wt, ΔgpoA, ΔahpD*, and *ΔtpxD* strains in A549 pneumocytes (A), RAW 264.7 macrophages (B), and differentiated PLB-985 neutrophils (C). To reduce ROS production during bacterial infection, A549 and RAW 264.7 cells were treated with 5 mM and 10 mM NAC, respectively, while untreated cells served as controls. Additionally, PLB-985 neutrophils were infected with these strains alongside PLB-985 *nox2-KO* cells, which have diminished ROS production. Data in all panels represent the mean ± SEM from at least three independent experiments. Statistically significant differences were assessed using a two-tailed test and are marked as *P* < 0.05 (*), *P* < 0.01 (**), or *P* < 0.001 (***).

To determine whether inhibition of ROS production affect the intracellular survival of *S. pneumoniae*, not only in the *wt*, but also in the Δ*gpoA, ΔtpxD*, and *ΔahpD* mutants, we used the same infection model for the A549 pneumocytes and RAW 264.7 macrophages, but with a treatment during infection of 5 mM NAC for 3 h for A549 cells, and 10 mM NAC for 1 h for RAW 264.7 cells. This treatment reduced approximately 10 and 3 times the intracellular ROS levels in the A549 and RAW 264.7 cells, respectively, as described [8]. In both cells lines, the *wt* strain, as well as and the *ΔgpoA, ΔtpxD* and *ΔahpD* mutants, showed an increased intracellular survival caused by the low ROS levels after the NAC treatment (Fig. 5A-B, Fig S5A-B). The PLB-985 neutrophils were not treated with NAC, but we included in this assay the *gp91*^*phox*^ KO-PLB-985 cells (hereafter referred to as PLB-985-*nox2* KO) [22]. This is a stable cellular line derivative from PLB-985 that lacks the gp91^phox^ subunit of the NADPH oxidase 2 (Nox2), because the *gp91*^*phox*^ gene was knocked out [22], which present a reduction of 3 times in the intracellular ROS levels [8]. Both cell lines were differentiated into neutrophil-like granulocytes by DMSO treatment, which were checked by CD11b expression (80%) and the presence of dead cells measured by propidium iodide assays (2%), as described [8]. The *wt, ΔgpoA, ΔtpxD* and *ΔahpD* mutants grew significantly more in *nox2*-KO, which also present low ROS levels [22], than in the *wt* PLB-985 cells (Fig. 5C, Fig S5C). Taken together, these findings suggest that, regardless of the type of host cell, *S. pneumoniae* must tolerate the ROS levels produced by host cells. Furthermore, the intracellular survival mechanism of the bacterium requires the involvement of each enzyme comprising the H_2_O_2_ detoxification system, including GpoA, TpxD, and AhpD.

### 3.4 GpoA contributes to levofloxacin persistence under oxidative stress during intracellular infection in host cells

Previous studies in our laboratory have shown that oxidative stress conditions can promote persistence against fluoroquinolones, such as levofloxacin, in *S. pneumoniae* [3, 8, 25]. Given that GpoA plays a protective role against oxidative stress, we investigated whether deletion of the *gpoA* gene would impact on levofloxacin persistence. First, we assessed levofloxacin persistence in bacterial cultures by pre-exposing the *wt* and *ΔgpoA* strains to 20 mM H_2_O_2_ for 30 min, followed by treatment with 6 mg/ml levofloxacin for 5 h [8]. As expected, the *wt* strain exhibited increased persistence after oxidative stress induction, consistent with previous reports [8]. In contrast, the *ΔgpoA, ΔahpD and ΔtpxD* mutants displayed significantly reduced survival after levofloxacin exposure compared to the *wt* strain (Fig. 6A, Fig S6). We also constructed the *ΔgshT* mutant that lacks the glutathione transporter [9], and we found the same phenotype that the *ΔgpoA* mutant, even with the addition of glutathione. This suggests that ROS detoxification mediated by GpoA, as well as AhpD and TpxD, are important for the induction of antibiotic persistence under oxidative stress conditions, which is dependent of the imported glutathione (Fig. 6A; Figs. S6).

**Fig 6.**
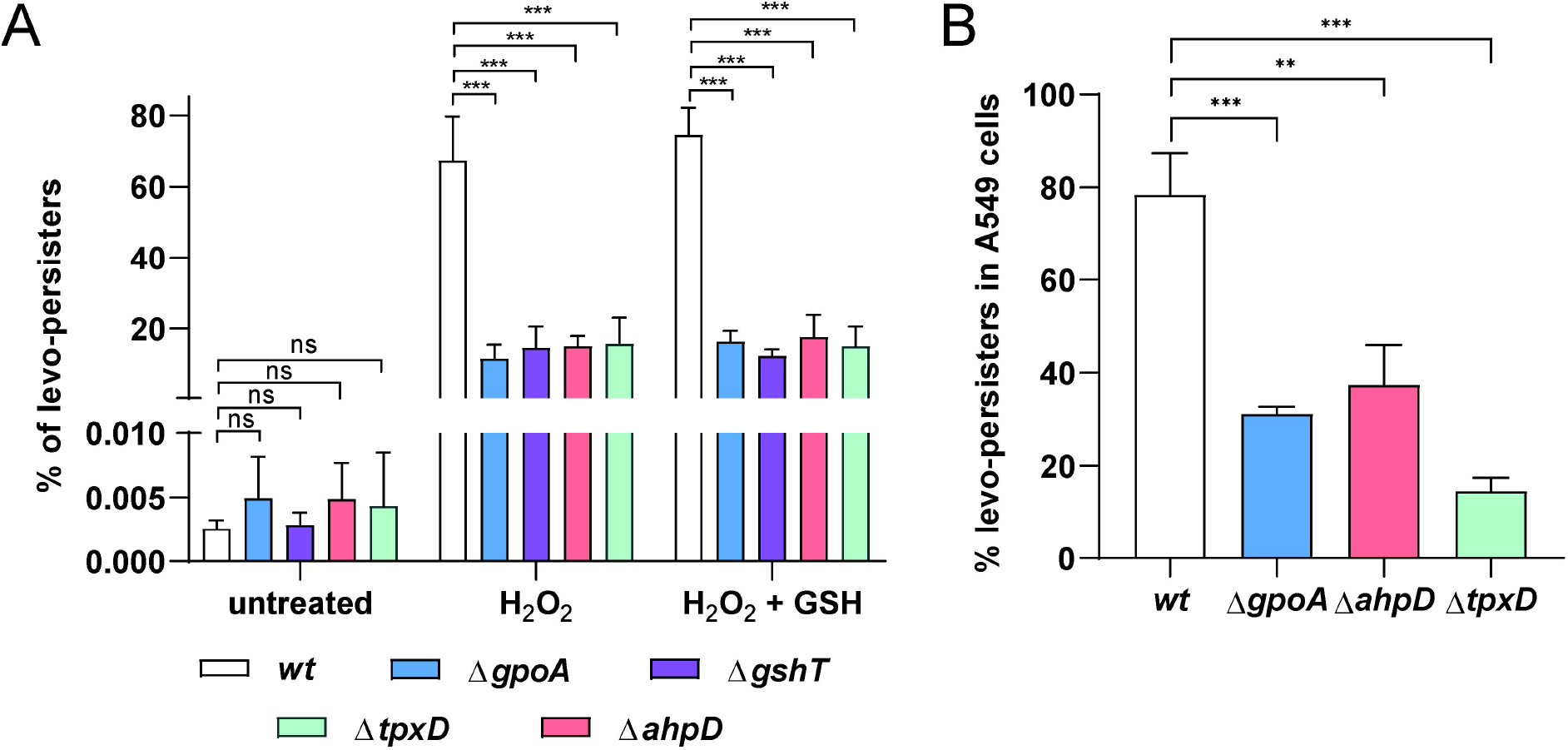
GpoA mediates levofloxacin persistence in bacterial cultures and during intracellular infection in A549 cells. (A) Levofloxacin persistence of the *wt, ΔgpoA, ΔgshT, ΔtpxD and ΔahpD* strains in the presence or absence of exogenous glutathione (GSH) in bacterial cultures. (B) Intracellular levofloxacin persistence was assessed in *wt* and Δ*gpoA* strains within A549 lung epithelial cells. Data in all panels represent the mean ± SEM of at least three replicates. Statistically significant differences were determined using a two-tailed test and are indicated as *P* < 0.01 (**), or *P* < 0.001 (***). Ns: not significant.

We next examined whether this phenomenon occurs during infection of host cells. The *wt, ΔgpoA, ΔahpD and ΔtpxD* strains were used to infect A549 pneumocytes. Following oxidative stress exposure within the host environment, the levofloxacin treatment revealed that the three mutants showed significantly lower survival rates in A549 cells compared to the *wt* strain, (Fig. 6B, Figs. S6). We also compared the phenotype of the *ΔgpoA* mutant in RAW 264.7 macrophages and PLB-985 neutrophils, obtaining consistent results compared with A549 cells (Fig. S7). These findings support a role for GpoA, along with AhpD and TpxD, in enabling *S. pneumoniae* to survive antibiotic treatment under oxidative stress, both in vitro and within the host.

### 3.5 GpoA is required for the pneumococcal tissue invasion in a mouse model infection

Our results referred to intracellular survival of *S. pneumoniae* in host cells (Fig. 3), in addition to the essentiality of glutathione that must be imported from host tissues via GshT [9], prompted us to study the putative impact of GpoA on the pneumococcal pathogenesis using an intranasal mouse model to mimic a natural pneumococcal infection [37]. We selected the murine BALB/c strain because it provides a more consistent response to infection, and is less vulnerable to pneumococcal infection compared to other mouse strains. This distinctive trait allows the animals to survive for more than 24 h and makes it possible to measure bacterial clearance in various tissues [37]. To determine the role of GpoA during pneumococcal infection, we constructed the *ΔgpoA* mutant in the capsulated D39 strain (serotype 2), which is virulent in BALB/c mice using intranasal inoculation [38], and its corresponding revertant strain (*wt gpoA*^+^). It is important to note that these mutants showed the same phenotype referred to H_2_O_2_ susceptibility (Fig. 3B) and intracellular survival (Fig. 5) that those displayed by the *ΔgpoA*, and *wt gpoA*^+^ strains constructed in the R801 genetic background (Fig. S8). Groups of BALB/c mice were challenged intranasally with 1×10^7^ CFU of the *wt, ΔgpoA*, and *wt gpoA*^+^ strains individually. After 24 h of inoculation, 100% of mice suffered moderated symptoms but survived to pneumococcal challenge. At that time, the animals were sacrificed to extract lung, liver, spleen and brain, and to determine the bacterial load from homogenates obtained from each organ. We observed that the *wt* strain showed a typical high bacterial load in lungs compared to other organs caused by the intranasal inoculation, as described [19] (Fig. 7A). The detection of infections in the spleen and liver (Fig. 7B-C) indicates that *S. pneumoniae* disseminated bacteremically to these organs within less than 24 hours following the onset of pneumonia. We observed that the bacterial load in brain is curiously high, suggesting that the pneumococcal infections could be reached by bacteremia but also through a nonhematogenous route [39, 40]. In all tissues, the *ΔgpoA* displayed a significant decrease in their bacterial loads, while the *wt gpoA*^+^ strain restored the *wt* infection levels (Fig. 7). These findings confirmed the relevance of GpoA in the pathogenesis of *S. pneumoniae* observed in both infection models, host cells and mice.

**Fig 7.**
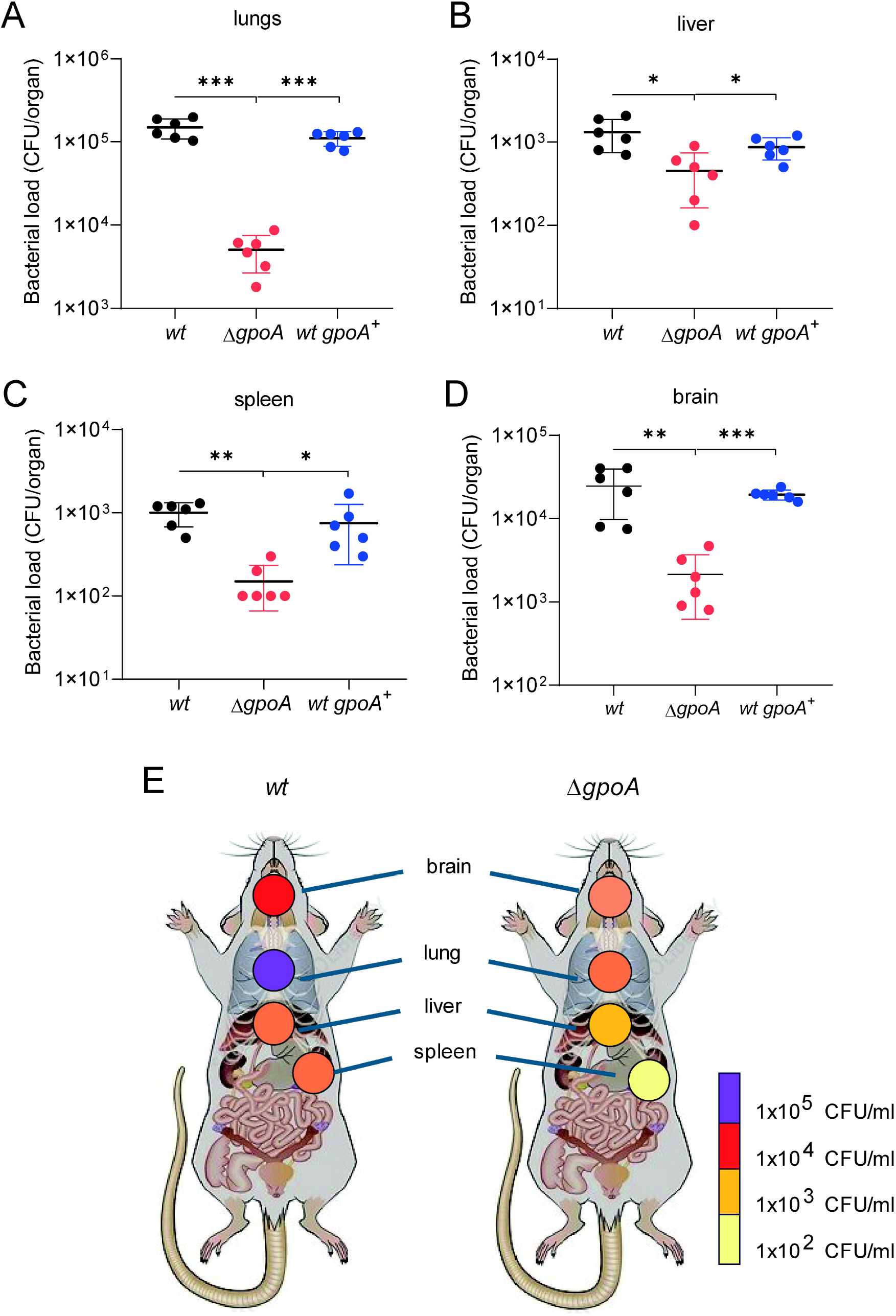
The role of GpoA in pneumococcal virulence was evaluated using a murine infection model. Panels A through D depict bacterial burdens quantified as CFU recovered from the lungs (A), livers (B), spleens (C), and brains (D) of mice infected with the virulent D39 *wt*, Δ*gpoA*, and *gpoA*+ (revertant) strains. Each panel presents the mean ± standard error of the mean (SEM) derived from a minimum of three independent experiments. Statistical significance was assessed via a two-tailed test, with differences denoted as *P* < 0.05 (*), *P* < 0.01 (**), and *P* < 0.001 (***). Panel E provides a schematic overview of the murine infection model, illustrating organ-specific bacterial loads through a color gradient corresponding to CFU.

## 4. Discussion

Glutathione (GSH) is recognized as the most abundant and essential endogenous antioxidant, functioning as the primary non-protein thiol constituent within the ROS detoxification system in living cells [41]. The genes responsible for GSH biosynthesis, which encodes for the γ-glutamylcysteine synthetase (GshA) and the glutathione synthetase (GshB), are conserved within Proteobacteria. Remarkably, *S. pneumoniae* lacks the capacity for endogenous GSH synthesis; however, this human pathogen possesses a GSH transporter that facilitates the uptake of GSH from host tissues [9], a phenomenon corroborated by the findings of the present study.

During the evolution of Cyanobacteria and Proteobacteria, GSH-dependent ROS detoxification systems incorporated enzymes that play crucial roles in these mechanisms and use GSH as a cofactor, such as GSH peroxidases, GSH reductases, glutaredoxins, and GSH-S-transferases [41]. From an evolutionary perspective, the presence of GpoA in *S. pneumoniae*, given its high homology with human and other streptococcal glutathione peroxidases (Fig. S1), suggests that this enzyme is highly conserved. In *S. pneumoniae*, it was likely acquired through horizontal gene transfer or evolutionary convergence and subsequently adapted to optimize resistance against oxidative attacks within the host.

In this work, we observed that transcripts encoding GpoA are significantly upregulated following exposure to H2O_2_, aligning with its specialized role in H_2_O_2_ detoxification. The deletion of the *gpoA* gene, as well as other associated genes including *tpxD* and *ahpD*, led to pronounced sensitivity to _2_0 mM H_2_O_2_, underscoring their involvement in ROS defense mechanisms. Notably, each individual mutant exhibited a comparable phenotype, characterized by an approximate 12-fold reduction in survival, while still maintaining a residual ability to endure H_2_O_2_ exposure. These findings indicate that GpoA, TpxD, and AhpD contribute significantly to the oxidative stress response and suggest that these three peroxidases act concurrently to form a robust detoxification system against H_2_O_2_.

While the primary focus of this investigation is the characterization of GpoA, the findings related to AhpD merit detailed consideration. Previous studies have reported that AhpD alkylhydroperoxidase exhibits weak thiol peroxidase activity [28] and is critical for pneumococcal virulence in murine models [29]. Furthermore, a double mutant lacking the operon encompassing both *ahpD* and *spr0371* genes demonstrated enhanced resistance to H_2_O_2_ concentrations of 9, 18, and 37 mM in the R6 strain [29]. This observation contrasts with the pronounced sensitivity to 20 mM H_2_O_2_ that we observed in the single *ΔahpD* mutant derived from the R801 strain (a derivative of R6) in bacterial cultures (Fig. 3B) and during intracellular survival within host cells (Fig. 5). We also analyzed this phenotype in the *ΔahpD* mutant constructed in the D39 genetic background (Fig. S3), which is the ancestral strain from which the R6 and R801 strains were derived. Notably, this susceptibility phenotype parallels those observed in the *ΔgpoA* and *ΔtpxD* mutants (Fig. 3B). The involvement of these peroxidases in the detoxification of H_2_O_2_ is further corroborated by the upregulation of their respective transcripts following exposure to 20 mM H_2_O_2_ (Fig. 3A) and the lower intracellular survival in host cells (Fig. 5). We propose that these phenotypic manifestations are more consistent with the attenuated virulence reported for the double *ahpD-spr0371* mutant in the D39 strain [29]. The discrepancies observed by the authors in H_2_O_2_ susceptibility assays in the R6 strain are more likely attributable to the *spr0371* deletion than to putative strain-specific differences.

Beyond molecular and functional analyses, biochemical assays confirmed that GpoA presents GSH-dependent activity analogous to that observed in eukaryotic glutathione peroxidases [12]. This evidence substantiates the proposition that GpoA operates within a H_2_O_2_ detoxification pathway in *S. pneumoniae*, thereby contributing to the mitigation of oxidative stress. The preservation of the active site architecture, notably the cysteine residue at position 36 (Cys^36^), coupled with the marked reduction in enzymatic activity exhibited by the GpoA^Cys36Ala^ variant relative to the wild-type enzyme, provides compelling biochemical support for the essential role of this residue in mediating GpoA’s peroxidase function.

The conserved cysteine residue at position 36 in GpoA corresponds to the selenocysteine (Sec) residue at position 46 in human GPX4 (Fig. 2). Selenocysteine residues are situated within the active sites of selenoproteins and exhibit superior redox catalytic efficiency relative to cysteine, which contains sulfur rather than selenium [35]. Recognized as the 21st naturally occurring amino acid, Sec is present across the domains Bacteria, Archaea, and Eukarya and is encoded by the UGA codon. Following this codon, selenoprotein mRNAs possess a selenocysteine insertion sequence (SECIS) element that mediates the incorporation of Sec in bacterial systems [36]. Interestingly, downstream of the cysteine-encoding UGU codon, a SECIS element was identified within the *gpoA* gene sequence (Fig. S4), indicating a potential substitution of Sec by Cys at this site during evolution. This amino acid replacement may maintain the critical redox functionality of the protein without necessitating the incorporation of selenocysteine. Previous studies have documented bacterial selenoproteins involved in redox metabolism, including a glutaredoxin in *Clostridium oremlandii* [42] and a thioredoxin in *Treponema denticola* [42]. The absence of Sec in GpoA likely represents an evolutionary adaptation that could affect the enzyme’s catalytic characteristics and regulatory mechanisms.

The functional relevance of GpoA was also evidenced through intracellular survival of *S. pneumoniae* in host cells. The deletion of the *gpoA* gene resulted in a marked decrease in bacterial survival across various host cell types characterized by differing levels of H_2_O_2_ production, including pneumocytes, macrophages, and neutrophils. This finding underscores the essential role of antioxidant defense mechanisms in facilitating bacterial adaptation to the intracellular milieu. Furthermore, the improved survival of the *gpoA*-deficient mutant under low ROS level, achieved via NAC treatment, further substantiates the critical role of oxidative stress resistance in *S. pneumoniae* survival during infection. One plausible explanation for these results is that the Δ*gpoA* mutant is unable to effectively counteract the elevated ROS concentrations encountered within the intracellular environment of host cells, despite the partial compensatory activity of the TpxD and AhpD peroxidases (Fig. 5) in the absence of GpoA.

In our laboratory, we have identified and characterized the first mechanism of antibiotic persistence in *S. pneumoniae*. This pathogen demonstrates the ability to survive exposure to lethal fluoroquinolone levels—including levofloxacin, moxifloxacin, and ciprofloxacin— following prior exposure to H_2_O_2_. This persistence phenomenon is evident not only in bacterial cultures but also during infections of host cells. The fluoroquinolone persistence mechanism is intricately linked to the oxidative stress response of *S. pneumoniae* and is dependent on the enzymatic activities of SodA and TpxD, which function to neutralize superoxide radicals and hydrogen peroxide, respectively [3, 8].

In this study, we demonstrated that the decreased FQ persistence observed in the *ΔgpoA* mutant parallels that reported for the Δ*tpxD* mutant [8] as well as the *ΔahpD* mutant (Fig. 6). Our findings provide a more comprehensive understanding of the FQ persistence mechanism, confirming that the H_2_O_2_ detoxification system—comprising the GpoA, TpxD and AhpD peroxidases—is critical for pneumococcal survival under lethal FQ concentrations. This requirement holds true in both intracellular environments and extracellular inflammatory sites characterized by elevated oxidative stress. Such conditions may facilitate the emergence of genetic resistance, underscoring the clinical significance of these findings for the management of pneumococcal infections [3, 8].

Previous studies have demonstrated the involvement of the oxidative stress response in pneumococcal virulence, specifically implicating TpxD [4] and SodA [6, 43], through the use of murine infection models. In the present study, the role of GpoA in virulence was further validated using a mouse infection model wherein animals were intranasally inoculated to replicate the natural route of human infection. Mice infected with the D39 *ΔgpoA* mutant strain exhibited a significant reduction in bacterial loads across multiple tissues, including the lung, brain, liver, and spleen, relative to those infected with the wild-type strain. These results suggest that GpoA is critical not only for resistance to oxidative stress but also for effective invasion and systemic dissemination during infection.

## 5. Conclusion

*S. pneumoniae* is traditionally considered a typical extracellular pathogen; however, recent studies have demonstrated its transient intracellular presence in pulmonary and immune cells, where defense against reactive oxygen species (ROS) is essential for its survival [3]. This study provides a comprehensive analysis of the role of GpoA, a crucial enzyme involved in the oxidative stress response of *S. pneumoniae*, and the most well-characterized bacterial glutathione peroxidase to date. Our results reveal that GpoA is a key component of the bacterium’s antioxidant defense system, which also includes the TpxD thiol peroxidase [4, 5] and the AhpD alkylhydroperoxidase [28, 29]. GpoA significantly enhances the bacterium’s ability to withstand hostile environments generated by the host immune response, and it also affects in vivo survival, as the *ΔgpoA* mutant exhibited reduced survival compared to the wild-type strain in various organs of infected mice. Furthermore, we extended our analysis to explore the association between oxidative stress response and antibiotic persistence, highlighting the clinical relevance of this enzyme as a potential factor complicating the treatment of *S. pneumoniae* infections.

## CRediT authorship contribution statement

**Victoria E. Zappia**: Writing – review & editing, Writing – original draft, Investigation, Formal analysis, Data curation.

**Mirelys Hernandez-Morfa**:Writing – review & editing, Investigation, Formal Analysis, Conceptualization.

**Luciana Raya-Plasencia**:Writing – review & editing, Investigation, Formal analysis, Conceptualization.

**Nadia B. Olivero**:Writing – review & editing, Investigation, Formal analysis, Conceptualization.

**Paulo R. Cortes**:Writing – review & editing, Investigation, Formal analysis, Conceptualization.

**Jose Echenique**:Writing – review & editing, Writing – original draft, Supervision, Project administration, Funding acquisition, Project Administration, Formal analysis, Data curation, Conceptualization.

## Funding

This work was supported by the National Agency of Scientific and Technological Promotion (ANPCYT; PICT 2018, to JE), and the Scientific and Technological Secretary of the National University of Cordoba (SECYT-UNC 2020, to JE). JE is member of the Research Career of CONICET. V.E.Z., M-H-M, and N.B.O. were recipients of CONICET Ph.D. fellowships. The funders had no participation in the study design, data collection, analysis, publication decision, or manuscript preparation.

## Declaration of competing interest

The authors report that there are no competing interests to declare.

## Acknowledgments

We thank Gabriela Furlan, Noelia Maldonado and Alejandra Romero (Cell Culture Facility, Luciana Reyna (Molecular Biology Facility), Paula Abadie and Santiago Boccardo (Cytometry Facility), Pilar Crespo (Microscopy Facility), Victoria Blanco and Raul Villarreal (Animal Care Facility), Sergio Oms, Ariel Frontera, Diego Luti, Walter Requena, Cecilia Noriega, and German Romero (BIOPROAL - Animal Production Facility) for their skillful technical assistance at the CIBICI-CONICET/ Department of Clinical Biochemistry, School of Chemistry Sciences, National University of Cordoba. Thanks to Dr. Mary Dinauer (Indiana University) for your kind gift of PLB-985 PLB-985-KO cells, and Dr. Analia Trevani and Dr. Florencia Sabbione (IMEX-CONICET) for technical information.

## Appendix A. Supplementary data

Supplementary data to this article can be found online at xxx.

## Data availability

Data will be made available on request.

## Notes

### Competing Interest Statement

The authors have declared no competing interest.

## References

[1] A. R. Narciso, R. Dookie, P. Nannapaneni, S. Normark, and B. Henriques-Normark, Streptococcus pneumoniae epidemiology, pathogenesis and control. Nat. Rev. Microbiol., 23 (2025) 256–271. 10.1038/s41579-024-01116-z.

[2] H. Sati, E. Carrara, A. Savoldi, P. Hansen, J. Garlasco, E. Campagnaro, S. Boccia, J. A. Castillo-Polo, E. Magrini, P. Garcia-Vello, E. Wool, V. Gigante, E. Duffy, A. Cassini, B. Huttner, P. R. Pardo, M. Naghavi, F. Mirzayev, M. Zignol, A. Cameron, E. Tacconelli, and W. H. O. B. P. P. L. A. Group., The WHO Bacterial Priority Pathogens List 2024: a prioritisation study to guide research, development, and public health strategies against antimicrobial resistance, Lancet. Infect. Dis. 25 (2025) 1033–1043. 10.1016/S1473-3099(25)00118-5.

[3] M. Hernandez-Morfa, N. B. Olivero, V. E. Zappia, G. E. Pinas, N. M. Reinoso-Vizcaino, M. B. Cian, M. Nunez-Fernandez, P. R. Cortes, and J. Echenique, The oxidative stress response of Streptococcus pneumoniae: its contribution to both extracellular and intracellular survival, Front. Microbiol., 14 (2023) e1269843. 10.3389/fmicb.2023.1269843.

[4] B. Hajaj, H. Yesilkaya, R. Benisty, M. David, P. W. Andrew, and N. Porat, Thiol peroxidase is an important component of Streptococcus pneumoniae in oxygenated environments, Infect. Immun., 80 (2012) 4333–43. 10.1128/IAI.00126-12.

[5] J. P. Lisher, H. T. Tsui, S. Ramos-Montanez, K. L. Hentchel, J. E. Martin, J. C. Trinidad, M. E. Winkler, and D. P. Giedroc, Biological and Chemical Adaptation to Endogenous Hydrogen Peroxide Production in Streptococcus pneumoniae D39, mSphere, 2 (2017) e291–16. 10.1128/mSphere.00291-16.

[6] H. Yesilkaya, A. Kadioglu, N. Gingles, J. E. Alexander, T. J. Mitchell, and P. W. Andrew, Role of manganese-containing superoxide dismutase in oxidative stress and virulence of Streptococcus pneumoniae, Infect. Immun., 68 (2000) 2819–2826. 10.1128/IAI.68.5.2819-2826.2000.

[7] N. M. Reinoso-Vizcaino, M. B. Cian, P. R. Cortes, N. B. Olivero, M. Hernandez-Morfa, G. E. Pinas, C. Badapanda, A. Rathore, D. R. Perez, and J. Echenique, The pneumococcal two-component system SirRH is linked to enhanced intracellular survival of Streptococcus pneumoniae in influenza-infected pulmonary cells, PLoS Pathog., 16 (2020) e1008761. 10.1371/journal.ppat.1008761.

[8] M. Hernandez-Morfa, N. M. Reinoso-Vizcaino, N. B. Olivero, V. E. Zappia, P. R. Cortes, A. Jaime, and J. Echenique, Host Cell Oxidative Stress Promotes Intracellular Fluoroquinolone Persisters of Streptococcus pneumoniae, Microbiol. Spectr., 10 (2022) e0436422. 10.1128/spectrum.04364-22.

[9] A. J. Potter, C. Trappetti, and J. C. Paton, Streptococcus pneumoniae uses glutathione to defend against oxidative stress and metal ion toxicity, J. Bacteriol., 194 (2012) 6248–54. 10.1128/JB.01393-12.

[10] S. D. Bathige, N. Umasuthan, G. I. Godahewa, W. S. Thulasitha, I. Whang, S. H. Won, C. Kim, and J. Lee, Two variants of selenium-dependent glutathione peroxidase from the disk abalone Haliotis discus discus: Molecular characterization and immune responses to bacterial and viral stresses, Fish Shellfish Immunol. 45, (2015) 648–55. 10.1016/j.fsi.2015.05.028.

[11] C. Duong, H. J. Seow, S. Bozinovski, P. J. Crack, G. P. Anderson, and R. Vlahos, Glutathione peroxidase-1 protects against cigarette smoke-induced lung inflammation in mice, Am. J. Physiol. Lung Cell Mol. Physiol.,299 (2010) L425–433, 10.1152/ajplung.00038.2010.

[12] R. Brigelius-Flohe and M. Maiorino, Glutathione peroxidases, Biochim. Biophys. Acta, 1830 (2013) 3289–303. 10.1016/j.bbagen.2012.11.020.

[13] A. Brenot, K. Y. King, B. Janowiak, O. Griffith, and M. G. Caparon, Contribution of glutathione peroxidase to the virulence of Streptococcus pyogenes, Infect. Immun., 72 (2004) 408–13. 10.1128/IAI.72.1.408-413.2004.

[14] Y. Zhang, Q. Guo, X. Fang, M. Yuan, W. Hu, X. Liang, J. Liu, Y. Yang, and C. Fang, Destroying glutathione peroxidase improves the oxidative stress resistance and pathogenicity of Listeria monocytogenes, Front. Microbiol., 14 (2023) 1122623. 10.3389/fmicb.2023.1122623.

[15] F. Ursini, M. Maiorino, R. Brigelius-Flohe, K. D. Aumann, A. Roveri, D. Schomburg, and L. Flohe, Diversity of glutathione peroxidases, Methods Enzymol., 252 (1995) 38–53. 10.1016/0076-6879(95)52007-4.

[16] M. Kutlu and F. Susuz, Biochemical properties of glutathione peroxidase in Gammarus pulex, Bull. Environ. Contam. Toxicol., 73 (2004) 432–436. 10.1007/s00128-0040447-4.

[17] J. R. Echenique and M. C. Trombe, “Competence repression under oxygen limitation through the two-component MicAB signal-transducing system in Streptococcus pneumoniae and involvement of the PAS domain of MicB, J. Bacteriol., 183 (2001) 4599–608. 10.1128/JB.183.15.4599-4608.2001.

[18] P. R. Cortes, G. E. Pinas, A. G. Albarracin Orio, and J. R. Echenique, Subinhibitory concentrations of penicillin increase the mutation rate to optochin resistance in Streptococcus pneumoniae, J. Antimicrob. Chemother., 62 (2008) 973–977. 10.1093/jac/dkn322.

[19] P. R. Cortes, A. G. Orio, M. Regueira, G. E. Pinas, and J. Echenique, Characterization of in vitro-generated and clinical optochin-resistant strains of Streptococcus pneumoniae isolated from Argentina, J. Clin. Microbiol., 46 (2008) 1930–1934. 10.1128/JCM.02318-07.

[20] A. G. Albarracin-Orio, P. R. Cortes, M. Tregnaghi, G. E. Pinas, G. Argentinean Network Pneumococcus Study, and J. R. Echenique, A new serotype 14 variant of the pneumococcal Spain9V-3 international clone detected in the central region of Argentina, J. Med. Microbiol., 57 (2008) 992–999. 10.1099/jmm0.2008/000505-0.

[21] C.L.S.I., CLSI M100-Performance Standars for Antimicrobial Susceptibility Testing, 34th edition, CLSI, U.S., 2024.

[22] L. Zhen, A. A. King, Y. Xiao, S. J. Chanock, S. H. Orkin, and M. C. Dinauer, Gene targeting of X chromosome-linked chronic granulomatous disease locus in a human myeloid leukemia cell line and rescue by expression of recombinant gp91phox, Proc. Natl. Acad. Sci. U.S.A., 90 (1993) 9832–9836. 10.1073/pnas.90.21.9832.

[23] A. Panday, M. K. Sahoo, D. Osorio, and S. Batra, NADPH oxidases: an overview from structure to innate immunity-associated pathologies, Cell. Mol. Immunol., 12 (2015) 5–23. 10.1038/cmi.2014.89.

[24] M. H. Shahhosseiny, Z. Hosseiny, H. R. Khoramkhorshid, S. Azari, and M. A. Shokrgozar, Rapid and sensitive detection of Mollicutes in cell culture by polymerase chain reaction, J. Basic Microbiol., 50 (2010) 171–178. 10.1002/jobm.200800174.

[25] M. Hernandez-Morfa, N. M. Reinoso-Vizcaino, V. E. Zappia, N. B. Olivero, P. R. Cortes, C. C. Stempin, D. R. Perez, and J. Echenique, Intracellular Streptococcus pneumoniae develops enhanced fluoroquinolone persistence during influenza A coinfection, Front. Microbiol., 15 (2024) 1423995. 10.3389/fmicb.2024.1423995.

[26] G. E. Pinas, N. M. Reinoso-Vizcaino, N. Y. Yandar Barahona, P. R. Cortes, R. Duran, C. Badapanda, A. Rathore, D. R. Bichara, M. B. Cian, N. B. Olivero, D. R. Perez, and J. Echenique, Crosstalk between the serine/threonine kinase StkP and the response regulator ComE controls the stress response and intracellular survival of Streptococcus pneumoniae, PLoS Pathog., 14 (2018) e1007118. 10.1371/journal.ppat.1007118.

[27] J. A. Lanie, W. L. Ng, K. M. Kazmierczak, T. M. Andrzejewski, T. M. Davidsen, K. J. Wayne, H. Tettelin, J. I. Glass, and M. E. Winkler, Genome sequence of Avery’s virulent serotype 2 strain D39 of Streptococcus pneumoniae and comparison with that of unencapsulated laboratory strain R6, J. Bacteriol., 189 (2007) 38–51. 10.1128/JB.01148-06.

[28] Y. Meng, C. R. Sheen, N. J. Magon, M. B. Hampton, and R. C. J. Dobson, Structure-function analyses of alkylhydroperoxidase D from Streptococcus pneumoniae reveal an unusual three-cysteine active site architecture, J. Biol. Chem., 295 (2020) 2984–2999. 10.1074/jbc.RA119.012226.

[29] G. K. Paterson, C. E. Blue, and T. J. Mitchell, An operon in Streptococcus pneumoniae containing a putative alkylhydroperoxidase D homologue contributes to virulence and the response to oxidative stress, Microb. Pathog., 40 (2006) 152–160. 10.1016/j.micpath.2005.12.003.

[30] D. Moosmayer, A. Hilpmann, J. Hoffmann, L. Schnirch, K. Zimmermann, V. Badock, L. Furst, J. K. Eaton, V. S. Viswanathan, S. L. Schreiber, S. Gradl, and R. C. Hillig, Crystal structures of the selenoprotein glutathione peroxidase 4 in its apo form and in complex with the covalently bound inhibitor ML162, Acta Crystallogr. D Struct. Biol., 77 (2021) 237–248. 10.1107/S2059798320016125

[31] J. C. Lefevre, J. P. Claverys, and A. M. Sicard, Donor deoxyribonucleic acid length and marker effect in pneumococcal transformation, J. Bacteriol., 138 (1979) 80–86. 10.1128/jb.138.1.80-86.1979.

[32] C. D. Pericone, S. Park, J. A. Imlay, and J. N. Weiser, Factors contributing to hydrogen peroxide resistance in Streptococcus pneumoniae include pyruvate oxidase (SpxB) and avoidance of the toxic effects of the fenton reaction, J. Bacteriol., 185 (2003) 6815–6825. 10.1128/JB.185.23.6815-6825.2003.

[33] B. Spellerberg, D. R. Cundell, J. Sandros, B. J. Pearce, I. Idanpaan-Heikkila, C. Rosenow, and H. R. Masure, Pyruvate oxidase, as a determinant of virulence in Streptococcus pneumoniae, Mol. Microbiol., 19 (1996) 803–13. 10.1046/j.1365-2958.1996.425954.x.

[34] F. A. Arenas, W. A. Diaz, C. A. Leal, J. M. Perez-Donoso, J. A. Imlay, and C. C. Vasquez, The Escherichia coli btuE gene, encodes a glutathione peroxidase that is induced under oxidative stress conditions, Biochem. Biophys. Res. Commun., 398 (2010) 690–694. 10.1016/j.bbrc.2010.07.002.

[35] A. Borchert, J. Kalms, S. R. Roth, M. Rademacher, A. Schmidt, H. G. Holzhutter, H. Kuhn, and P. Scheerer, Crystal structure and functional characterization of selenocysteine-containing glutathione peroxidase 4 suggests an alternative mechanism of peroxide reduction, Biochim. Biophys. Acta Mol. Cell. Biol. Lipids, 1863 (2018) 1095–1107. 10.1016/j.bbalip.2018.06.006.

[36] S. Ishida, A. Gundlach, C. W. Kosonocky, and A. D. Ellington, Directed Evolution of a SelB Variant that Does Not Require a Selenocysteine Insertion Sequence Element for Function, ACS Synth. Biol., 14 (2025) 2681–2689. 10.1021/acssynbio.5c00106.

[37] N. Borsa, M. D. Pasquale, and M. I. Restrepo, Animal Models of Pneumococcal pneumonia, Int. J. Mol. Sci., 20 (2019) 4220. 10.3390/ijms20174220.

[38] L. C. Jacques, S. Panagiotou, M. Baltazar, M. Senghore, S. Khandaker, R. Xu, L. Bricio-Moreno, M. Yang, C. G. Dowson, D. B. Everett, D. R. Neill, and A. Kadioglu, Increased pathogenicity of pneumococcal serotype 1 is driven by rapid autolysis and release of pneumolysin, Nat. Commun., 11 (2020) 1892. 10.1038/s41467-020-15751-6.

[39] T. Audshasai, J. A. Coles, S. Panagiotou, S. Khandaker, H. E. Scales, M. Kjos, M. Baltazar, J. Vignau, J. M. Brewer, A. Kadioglu, and M. Yang, Streptococcus pneumoniae Rapidly Translocate from the Nasopharynx through the Cribriform Plate to Invade the Outer Meninges, mBio, 13 (2022) e0102422. 10.1128/mbio.01024-22.

[40] A. Marra and D. Brigham, Streptococcus pneumoniae causes experimental meningitis following intranasal and otitis media infections via a nonhematogenous route, Infect. Immun., 69 (2001) 7318–7325.10.1128/IAI.69.12.7318-7325.2001.

[41] C. Cassier-Chauvat, F. Marceau, S. Farci, S. Ouchane, and F. Chauvat, The Glutathione System: A Journey from Cyanobacteria to Higher Eukaryotes, Antioxidants (Basel), 12 (2023) 1199. 10.3390/antiox12061199.

[42] M. J. Kim, B. C. Lee, J. Jeong, K. J. Lee, K. Y. Hwang, V. N. Gladyshev, and H. Y. Kim, Tandem use of selenocysteine: adaptation of a selenoprotein glutaredoxin for reduction of selenoprotein methionine sulfoxide reductase, Mol. Microbiol., 79 (2011) 1194–1203. 10.1111/j.1365-2958.2010.07500.x.

[43] L. K. Mahdi, M. B. Van der Hoek, E. Ebrahimie, J. C. Paton, and A. D. Ogunniyi, Characterization of Pneumococcal Genes Involved in Bloodstream Invasion in a Mouse Model, PLoS One, 10 (2015) e0141816. 10.1371/journal.pone.0141816.

