## Supplementary material for "GpoA Glutathione Peroxidase Links Oxidative Stress Response, Antibiotic Persistence, and Virulence in *Streptococcus pneumoniae*": Table S1

**Table S1.** Bacterial strains and primers used in this work

| Name | Relevant characteristics | References |
|---|---|---|
| **Strains** | | |
| R801 | Derivative of R6 (unencapsulated derivative of D39) | 1 |
| R806 | R801, but *rpsL1* obtained by transformation of PCR product amplified by FrpsL and RrpsL primers from the CP1296 chromosomal DNA; $Sm^R$ | 2 |
| D39 | Wild-type, highly virulent Serotype 2 strain | 1 |
| Δ*ahpD* | R806, but Δ*ahpD::kan-rpsL⁺*, obtained by Janus cassette system (amplified from CP1296 chromosomal DNA) and pneumococcal DNA amplified with the FahpDA1, RahpDA1, FahpD2 and RahpD2 primers. $Km^R$, $Sm^S$ | This work |
| Δ*gpoA* | R806, but Δ*gpoA:: kan-rpsL⁺*, obtained by Janus cassette system (amplified from CP1296 chromosomal DNA) and pneumococcal DNA amplified with the FgpoAA1, RgpoAA1, FgpoA2 and RgpoA2 primers. $Km^R$, $Sm^S$ | This work |
| *gpoA*$^{C36A}$ | R801, obtained by PCR-based site-directed mutagenesis. Cysteine-to-Alanine substitution at position 36 of GpoA (*gpoA*$^{C36A}$). $Km^S$, $Sm^S$ | This work |
| Δ*gshT* | R806, but Δ*gshT::kan-rpsL⁺*, obtained by Janus cassette system (amplified from CP1296 chromosomal DNA) and pneumococcal DNA amplified with the FgshTA1, RgshTA1, FgshT2 and RgshT2 primers. $Km^R$, $Sm^S$ | This work |
| Δ*sodA* | R806, but Δ*sodA::kan-rpsL⁺*, obtained by Janus cassette system (amplified from CP1296 chromosomal DNA) and pneumococcal DNA amplified with the FsodA1, RsodA1, FsodA2 and RsodA2 primers. KmR, SmS. | 3 |
| Δ*spxB* | R806, but Δ*spxB::kan-rpsL⁺*, obtained by Janus cassette system (amplified from CP1296 chromosomal DNA) and pneumococcal DNA amplified with the FspxBA1, RspxBA1, FspxB2 and RspxB2 primers. KmR, SmS | 3 |
| Δ*tpxD* | R806, but Δ*tpxD*::*kan-rpsL⁺*, obtained by Janus cassette system (amplified from CP1296 chromosomal DNA) and pneumococcal DNA amplified with the FtpxDA1, RtpxDA1, FtpxD2 and RtpxD2 primers. KmR, SmS | 3 |
| Δ*gpoA* | D39, obtained by Janus cassette system (amplified from CP1296 chromosomal DNA) and pneumococcal DNA amplified with the FgpoAA1, RgpoAA1, FgpoA2 and RgpoA2 primers. KmR, $Sm^S$ | This work |
| *gpoA⁺* | D39, wild-type gpoA gene complemented. Obtained by counter-selection (using streptomycin) following transformation of the Δ*gpoA::kan-rpsL*+ strain with wild-type *gpoA* DNA. Acts as the parental strain control for *gpoA* mutants. $Km^S$, $Sm^S$ | This work |

| Primers | DNA sequences (5'-3') | Amplified gene |
|---|---|---|
| FJanus2 | TTGGATCCGCTAGCCTCGAGAAGCTTGGAACAAGTTATTACTTGAA GATGTCAG | Janus cassette |
| RJanus2 | AAGTCGACATCGATAGATCTTCTAGACCCTTTCCTTATGCTTTTGGAC | Janus cassette |
| FgpoA1 | ATATCAGGATCCAAGCTTAAAATCCTCTGTCACAAAGAGTTCCC | *gpoA* |
| RgpoA1 | CCGTATCGTCTCCAAAGATAAGCCCTCAATATGATAGAAACCG | *gpoA* |
| FgpoA2 | CCGTATCGTCTCCACGGTCACAATCTCACTATGATTAGGTTTCC | *gpoA* |
| RgpoA2 | GTTGGGGGATTATGACGAAGAGC | *gpoA* |
| FahpD1 | GGTGGAGCTGAAGTCATCAAACC | *ahpD* |
| RahpD1 | CCGTATCGTCTCCACGGAGCTTTATTCTCTCATAAGATTTTCGTTGT | *ahpD* |
| FahpD2 | GGTGGAGCTGAAGTCATCAAACC | *ahpD* |
| RahpD2 | GGTGGAGCTGAAGTCATCAAACC | *ahpD* |
| FgshT1 | GGTGGAGCTGAAGTCATCAAACC | *gshT* |
| RgshT1 | CCGTATCGTCTCCACGGCTTAACAGAAAAAGAGCATTTACGCC | *gshT* |
| FgshT2 | CCGTATCGTCTCCAAAGCCATTCTGAGTAAGGTGGGTGG | *gshT* |
| RgshT2 | GAAGTTGTCGACGGATCCCCATTCTGAGTAAGGTGGGTGG | *gshT* |
| FgpoA-C36 | CGTGGCAAAGTTCTCTTGATTGTCAACACTGCTACTGGTGCAGGTTTAACGCCC | *gpoA* |
| RgpoA | TTATAGTAGAGTTTGTATCGCCTCTTC | *gpoA* |

### References

1) Lefevre JC, Claverys JP, Sicard AM. Donor deoxyribonucleic acid length and marker effect in pneumococcal transformation. 1979. *J. Bacteriol*. 138:80-6.

2) Reinoso-Vizcaino NM, et al. The pneumococcal two-component system SirRH is linked to enhanced intracellular survival of *Streptococcus pneumoniae* in influenza-infected pulmonary cells. *PLoS Pathog* 16, e1008761 (2020).

3) Hernandez-Morfa, M., et al. Host cell oxidative stress promotes intracellular fluoroquinolone persisters of *Streptococcus pneumoniae*. *Microbiol Spectr* 10: e0436422. 2022
