## Supplementary figures for "GpoA Glutathione Peroxidase Links Oxidative Stress Response, Antibiotic Persistence, and Virulence in *Streptococcus pneumoniae*"

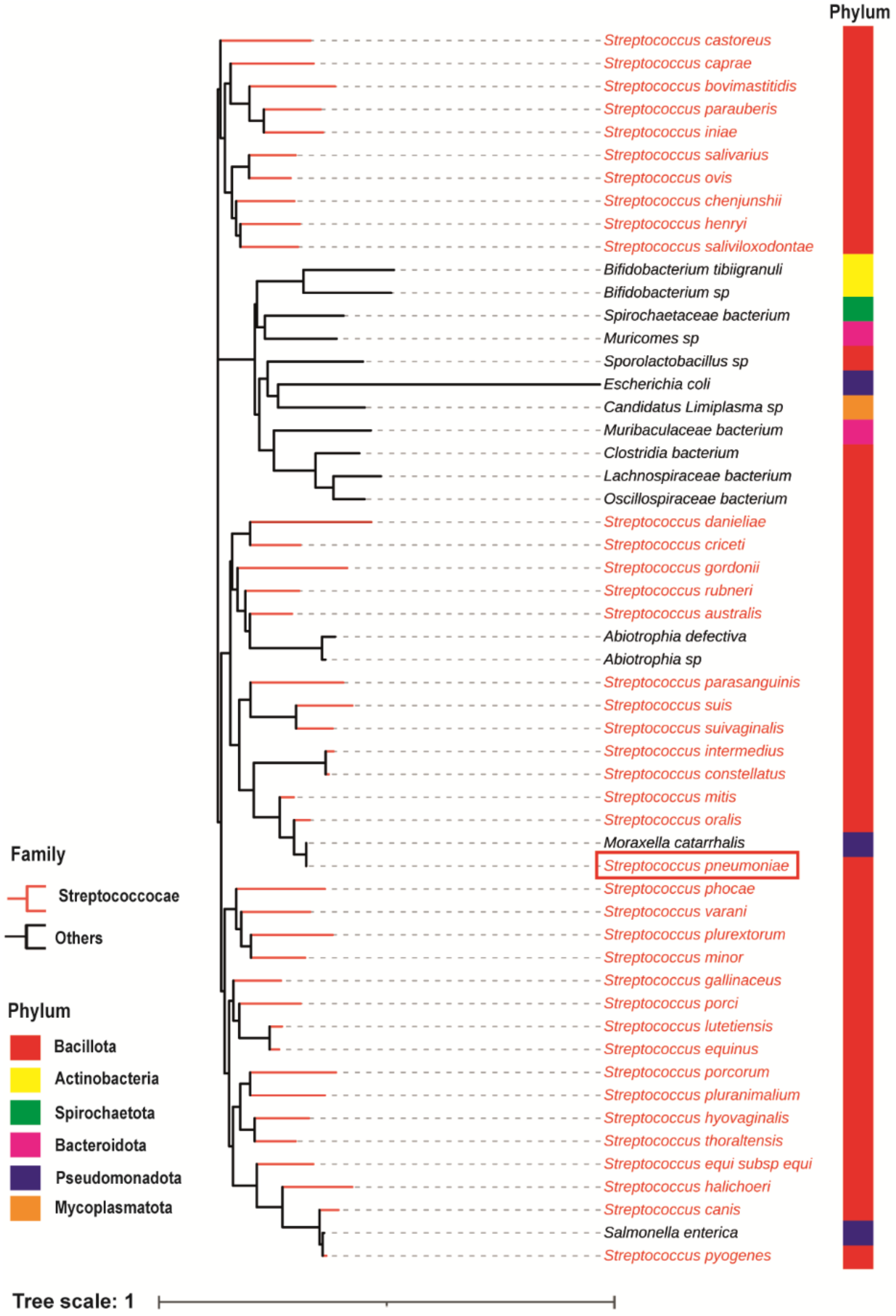


**Fig. S1. Phylogenetic tree of glutathione peroxidase (GpoA) proteins from representative bacterial phyla.** The tree was constructed using amino acid sequences of GpoA from multiple bacterial species to examine the evolutionary relationships and conservation of this enzyme across different phyla. Branches cluster according to the phylogenetic divergence of the sequences. The *Streptococcus pneumoniae* GpoA sequence is highlighted with a red box for easy identification.

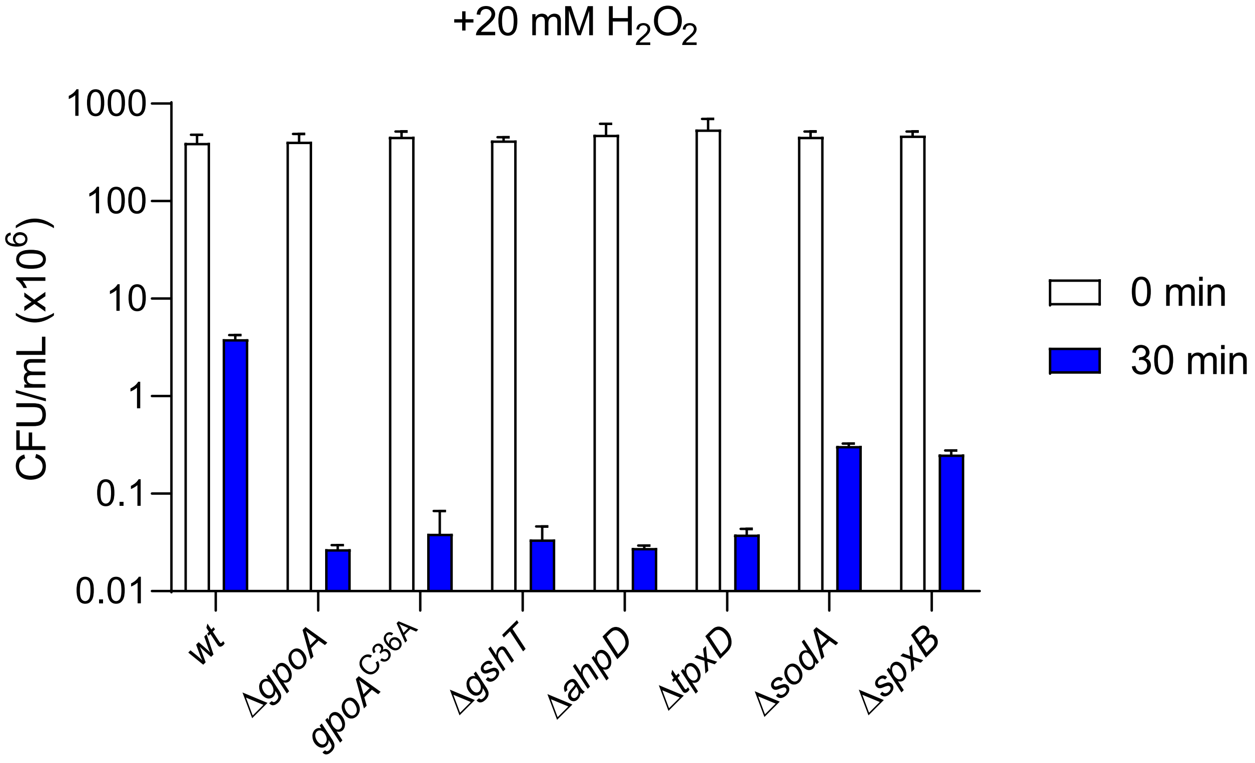


**Fig. S2. $H_2O_2$-susceptibility assays for different pneumococcal strains.** Colony-forming units (CFU) of the *wt*, Δ*gpoA, gpoA*$^{C36A}$*,* Δ*gshT,* Δ*ahpD,* Δ*tpxD,* Δ*sodA*, and Δ*spxB* strains were quantified after treatment with $H_2O_2$. Bacteria were grown in BHI broth to an $OD_{600nm}$ of 0.3 and then exposed to 20 mM $H_2O_2$ for 30 minutes. Then, bacterial cells were collected by centrifugation, washed with PBS, resuspended, and plated on blood agar to determine bacterial survival after 24 h. The corresponding percentage values (treated vs non-treated pneumococci) are shown in Figure 3B.

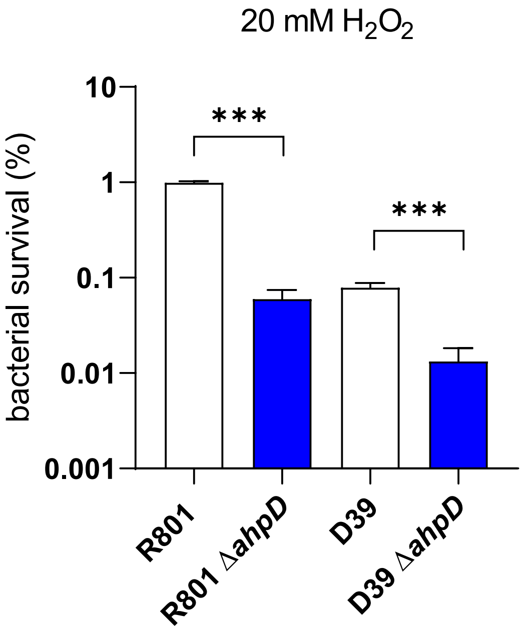


**Fig, S3. $H_2O_2$ susceptibility assays for the Δ*ahpD* mutant in the R801 and D39 genetic background.** Bacterial cells were exposed to 20 mM $H_2O_2$ for 30 min. Then, pneumococci were washed with PBS and plated on blood agar to estimate CFU/ml. The percentages shown in this panel were calculated from the CFU/ml values corresponding to these assays. Data represent the mean ± SEM of at least three replicates. Statistically significant differences were determined using a two-tailed test and are indicated as $P$ < < 0.001 (***),

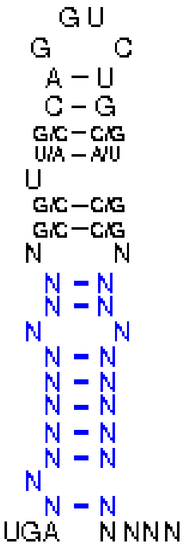


| | | |
|---|---|---|
| **Wanted Structure:** | [,[[[[[{[[/((.((((....))))))\]]}]]]]]].... | Prob.: |
| **mRNA-Sequence without optimizing the stability of the structure:** | GGCCUAACGCCCCUUGCAGGUCUGCAGGAGCUUUACGAAAGA | |
| | ..(.(((.((.((.((((....)))))).)).))).)..... | (0.01) |
| | (((.....))).((((((....))))))........(....). | (0.20) |
| **mRNA-Sequence after optimizing the stability of the structure:** | GGCUCCACGCCCCUUGCAGGUCUGCAGGAGCUUGGUGAAAGA | |
| | ..(.(((.((.((.((((....)))))).)).))).)..... | (0.24) |
| | ..(.(((.((.(((.(((....)))))).)).))).)..... | (0.24) |

**Fig. S4. Detection of a SECIS element into the *gpoA* sequence of *S. pneumoniae*.** (A) Loop constituted by the SECIS sequence, (B) mRNA sequence with structure and its probability for the SECIS-element region after UGA stop codon,

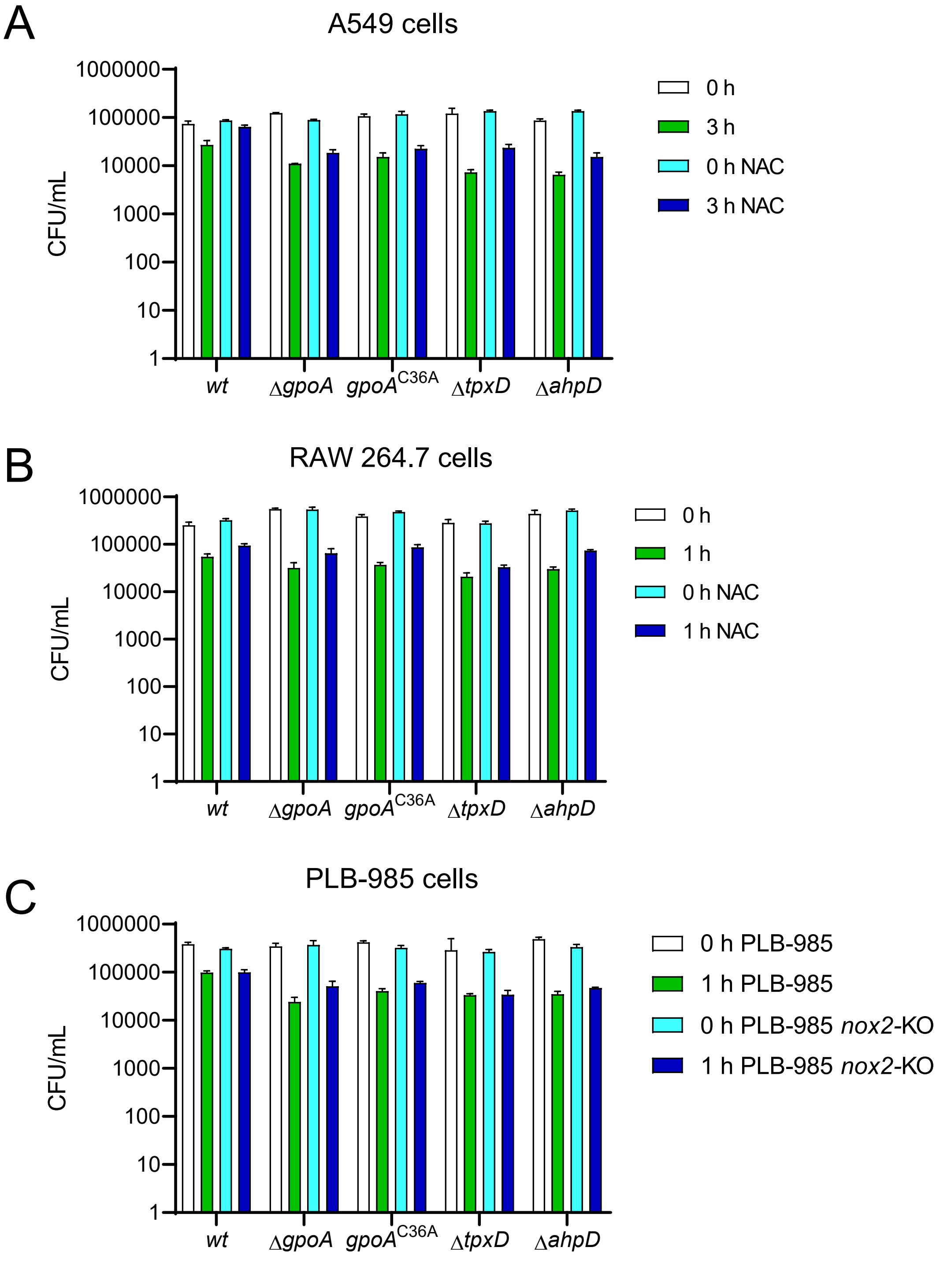


**Fig. S5. Intracellular survival assay of pneumococcal strains in different host cells.** A549 pneumocytes (A), Raw 264.7 macrophages (B), and differentiated PLB-985 or *PLB-985 nox2-KO* neutrophils (C) were cultured in DMEM supplemented with 10% FBS and 1% penicillin-streptomycin at 37°C in 5% $CO_2$. A549 and Raw 264.7 cells were pre-treated with 5 mM and 10 mM NAC, respectively; untreated cells served as controls. All cell types were infected with different strains at an MOI of 30:1. Intracellular survival was assessed using a gentamicin protection assay. At cell-specific time points, samples were collected, centrifuged to lyse host cells, and plated on blood agar for 16 h at 37°C to determine CFU. The CFU values were used to calculate the intracellular survival percentage shown in Fig. 5 for the *wt, ΔgpoA, gpoAC36A, ΔtpxD,* and *ΔahpD* strains.

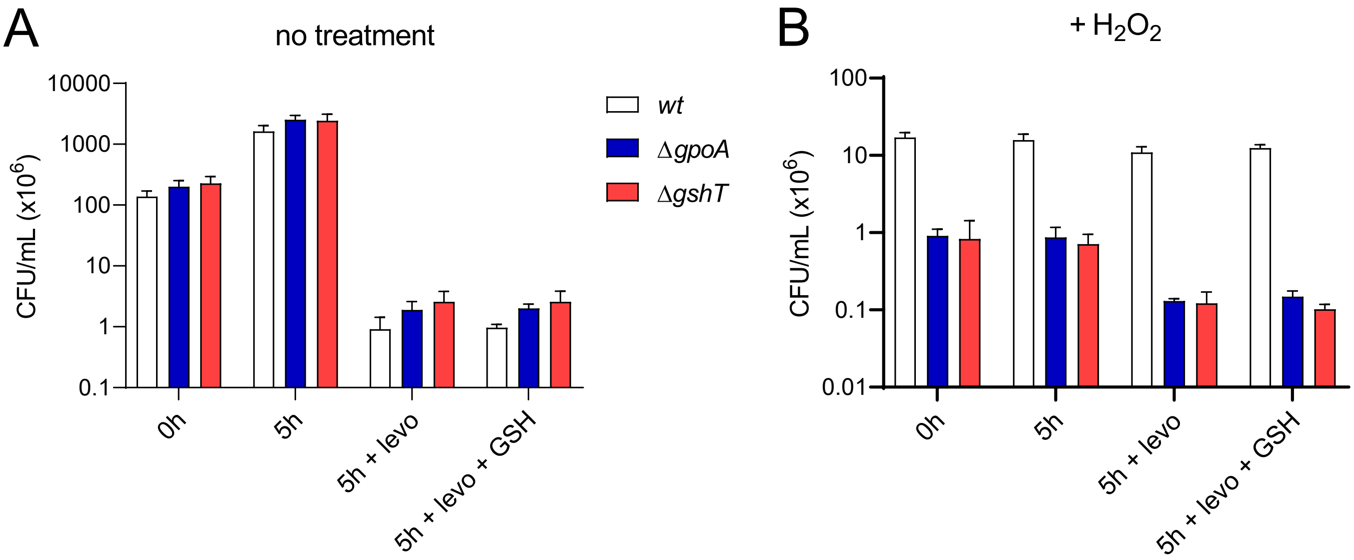


**Fig. S6. Fluoroquinolone persistence of different pneumococcal strains in bacterial cultures.** To assess FQ persistence, the *wt, ΔgpoA,* and *ΔgshT* strains were grown in BHI to mid-log phase without treatment (A) or exposed to 20 mM $H_2O_2$ for 30 minutes (B). Bacterial cells were collected by centrifugation, resuspended in BHI, and then exposed to 6 µg/ml levofloxacin and GSH for 5 h. The number of viable cells was measured as described in Fig. S1. These CFU/ml values were used to calculate the percentage of levo-persisters shown in Fig. 6A.

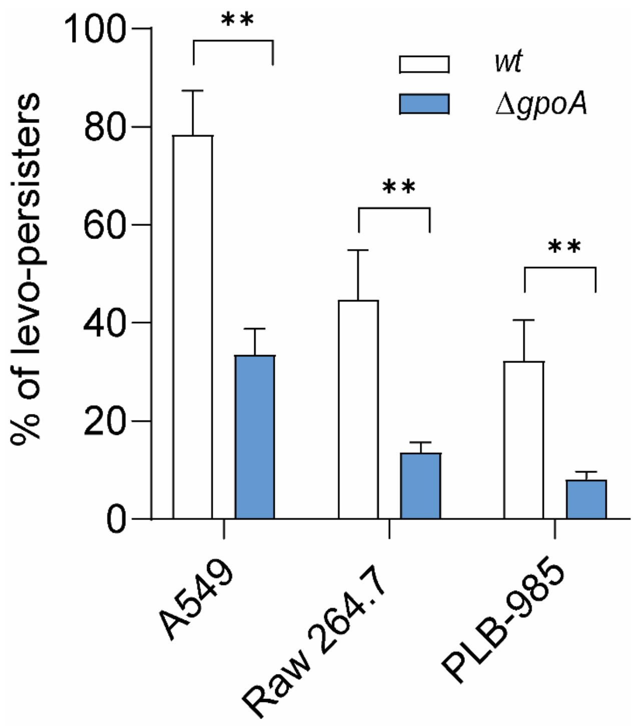


**Fig. S7. GpoA mediates levofloxacin persistence during intracellular infection in host cells.** Intracellular levofloxacin persistence was assessed in *wt* and Δ*gpoA* strains within A549 lung epithelial cells, Raw 264.7 macrophages, and differentiated PLB-985 neutrophils. Data in all panels represent the mean ± SEM of at least three replicates. Statistically significant differences were determined using a two-tailed test and are indicated as $P < 0.01$ (**),

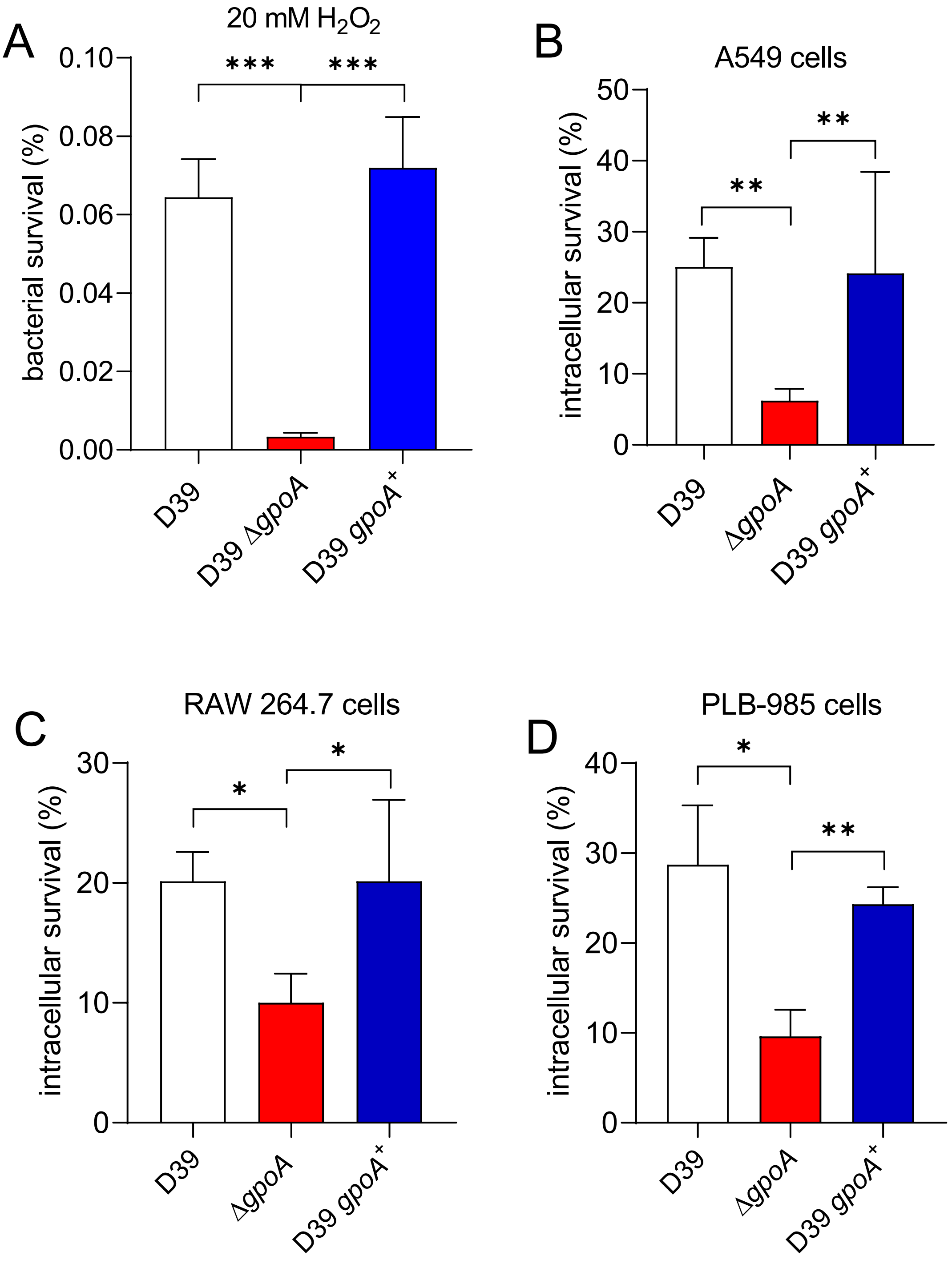


**Fig. S8. $H_2O_2$-susceptibility and intracellular survival assays of the *wt*, Δ*gpoA* and *gpoA*$^+$ *D39* strains*.*** Four-panel chart showing bacterial survival after $H_2O_2$ treatment (A), and intracellular survival in A549 pneumocytes (B), Raw 264.7 macrophages (C), and PLB-985 neutrophils (D) of the capsulated *wt*, Δ*gpoA*, and the *gpoA*$^+$ (revertant) D39 strains (serotype 2; virulent for mice). These assays were performed as described previously,
